# Vocal encoding of heritable sociability prevails over emotions in sheep bleats

**DOI:** 10.1101/2025.06.28.662115

**Authors:** Avelyne S. Villain, Alain Boissy, Gaetan Bonnafé, Christian Durand, Sébastien Douls, Paul Renaud-Goud, Marie-Madeleine Mialon, Dominique Hazard, Elodie F. Briefer

## Abstract

Vocalisations of animals are good indicators of their emotions. Temperament is known to influence the regulation and expression of emotions. However, how animal temperament affects their vocalisations and particularly their vocal expressions of emotions remains largely unexplored. Sociability is often measured as the behavioural reactivity to social separation and is a temperament trait intrinsically linked to emotional reactivity. Most social species respond to this challenging situation using contact calls. Here, we investigated whether the acoustic structure of these calls reflect sociability, emotions or both. Among 152 recorded female lambs, 42 belonged to two diverging sheep lines selected for high or low sociability. High bleats were recorded both in isolation (social challenge, all lambs) then before receiving a food treat (non-social challenge, selected lambs) to investigate the link between vocalisations, emotions and heritable sociability. The acoustic features of isolation bleats differed between lines, but it was not the case for pre-food treat bleats. Surprisingly, the genetic selection index and social behaviour explained better the structure of isolation bleats than the arousal. Last, encoding of individuality in isolation bleats was impaired by the genetic selection. Our findings show that selecting for sociable animals affects vocal signatures in calls produced during a social challenge, leading us to hypothesize a socio-acoustic co-selection.

## 1. Introduction

Vocalisations encode both static (i.e. relatively stable) information about the sender, such as its sex, identity or social rank [1,2], as well as more dynamic information about its internal motivation and emotional states [3]. Indeed, variations in acoustic signaling in term of call types, rate of production of these types or spectro-temporal features of vocalisations arising as a function of the sender’s emotions have been discovered in many non-human animals and show construct validity as indicators of emotional states [3–5]. Regulation and expression of dynamic features such as emotions depend on more stable traits, namely personality, in humans [6,7], with some evidence suggesting that this is also the case in non-human animals [8]. As a result, a link between temperament or personality and vocal expressions of emotions is expected, but has rarely been explored in non-human animals.

Temperament traits and personalities are defined as consistent and stable individual differences in response to species-specific environmental or social stimuli [9]. Although personalities may be shaped by environmental factors, temperament traits represent the heritable core part of these individual differences. Consequently, selective breeding of divergent lines for opposite temperament traits provide experimental models to study phenotypic consequences of temperament in ecology [9,10], medical [11] or agricultural research [12]. Boldness-shyness, exploration-avoidance, activity, sociability, aggressiveness or emotional reactivity, are temperament traits commonly assessed in both humans and non-human animals, including farm animals [9,13–15]. Most of these traits are inherently linked to, and measured as, individual differences in physiological and behavioural responses to challenging situations [9–11], and are thus intrinsically linked to emotions [16,17]. In particular, sociability may be defined as “an individual’s tendency or propensity to associate with other individuals, where the association is not driven by reproduction or aggression” [18] and may be partly measured as “an individual’s reaction to the absence of conspecifics” [9], i.e. quantifying the stress response and/or exacerbated behaviours to isolation.

In response to challenging situations, many animal species respond using vocalisations (alarm calls, contact calls). Yet, studies on the link between the acoustic structure of vocalisations (i.e. variation in duration, frequencies and amplitude) and temperament traits are rare, despite their potential to highlight non-invasive and rapid indicators of temperament. A few studies have incorporated call rate in their behavioural assessment of temperament [16,19–22]. In some cases, call types were produced in a context-specific way [19,23], which may emphasize the fact that some call types are used in specific situations and hence convey information about these contexts. Going further into the acoustic structure of vocalisations produced during temperament assessment, Leliveld et al (2007) focused on two common call types emitted by pigs, grunts and squeals, to investigate vocal correlate of emotional reactivity [24]. This study showed that grunts could be consistently divided into high and low frequency grunts, and that high frequency grunts were indicative of more active, more explorative and less stressful individuals. As vocalisations are strongly affected by emotions [3–5], and the range of emotions experienced by an individual is driven by its temperament, we can predict that the acoustic structure of vocalisations may reflect the temperament of non-human animals in a broader range of species as currently demonstrated [24], similarly to vocal correlates of personality in human voice [25]. Within a social group, acoustic correlates of temperament could be very useful for receivers, as it would allow them to predict the behaviour of others.

Sheep (*Ovis aries)* are highly gregarious and vocal species [26]. Their vocal repertoire contains two main structural call types: high and low bleats, differentiated by the opening of the mouth during the call (open and closed, respectively). High and low bleats are context specific; namely, high bleats are usually produced in higher arousal states, especially in negative situations such as isolation from conspecifics [26], and low bleats in lower arousal states, such as during short distance mother-lamb communication [27,28]. The acoustic structure of sheep bleats encodes individuality from an early stage after birth and is maintained over time [27,29], hence enabling early vocal recognition, especially between mother and lamb [28,30] as well as between kin and non-kin lambs [31], which is at the basis of the strong social bonds on which the lamb’s survival depends. Several temperament traits, such as ‘exploration-avoidance’, ‘boldness-shyness’ or ‘sociability’, were identified in sheep [14,32], in addition to their associations with cognitive and physiological characteristics [14]. Vocal production rate (i.e. the number of vocalisation produced per time unit) was found as indicative of sociability [32]. In addition, selective breeding based on sociability revealed that the rate of calls emitted during isolation from conspecifics was heritable and that highly sociable sheep genetic line produced bleats at a higher rate than less sociable ones [33]. However, whether the acoustic structure of sheep bleats also encodes sociability remains unknown.

In the present study, we studied a population of sheep consisting of two divergent lines of female sheep selected for high or low sociability for over 11 years (at the time of data collection) [33]. We evaluated the consequences of three generations of selection for sociability on the structure of sheep bleats recorded in two contexts that trigger emotions of different valence (negative/unpleasant *versus* positive/pleasant) and vary in their social context (social challenge/isolation from conspecifics *versus* prior to receiving a food treat in their baseline social group), to investigate which factors among ones explained most variation in bleat structure and study potential vocal correlates of temperament in sheep. We designed two non-exclusive hypotheses to explain bleat acoustic structure: the emotional hypothesis and the socio-acoustic co-selection hypothesis. According to the emotional hypothesis, the isolation test should trigger a higher level of arousal for high-sociability sheep than low-sociability ones, which should explain variation in acoustic structure, while the ‘anticipation’ of a food treat should not induce differential arousal states between the two lines. Considering vocal indicators of negative arousal, which are largely consistent across species [3], this should result in bleats of longer duration and with higher frequencies in the isolation bleats of high-sociability sheep compared to low-sociability individuals. Based on the socio-acoustic co-selection hypothesis, selecting for sociability also select for acoustic characteristics of vocalisations. Hence, genetic selection for sociability and social behaviour should better explain variation in acoustic structure than emotional arousal per se (when both factors are considered). We first tested for an acoustic signature of the two divergent lines of sheep, in the two recorded contexts. We predicted that, if selection for sociability drives the acoustic structure of bleats, a ‘line acoustic signature’ should be stronger in calls produced in a context of social challenge (isolation) than in a context without a social challenge (food treat ‘anticipation’). We then tested the effect of several potential predictors (biometry, emotional arousal, selection indexes and social behaviour) on the acoustic structure of isolation bleats to disentangle the emotional from the socio-acoustic co-selection hypothesis. Last, as the salience of vocal individuality (i.e. how much acoustic structure differ between individuals) has been shown to be affected by both emotional arousal [34] and valence [35], we additionally evaluated the potential consequences of the divergent selection for sociability on individual signature in bleats, a key stone for social communication.

## 2. Materials and Methods

### 2.1 Ethical note

This study was approved by the Animal Ethics Committee SCIENCE ET SANTE ANIMALES N°115 of the Veterinary School of Toulouse under the number: SSA_2022_003.

### 2.2 Animals and management

The experimental animals were Romane breed sheep, reared at the INRAE experimental farm of La Fage (Saint-Jean Saint-Paul, Causse du Larzac, France) exclusively outdoors [36]. The flock comprises about 250 reproductive females reared on 280 ha of rangelands (see [37] for details about the farming system and management characteristics). This population has been established in 2000 and selections based on sociability and reactivity to human started in 2011. Therefore, the selection procedure and corresponding analyzes summarised below are conducted on a yearly basis for the purpose of the long-term population (see more details here [38]), while recording of vocalisations and corresponding analyzes were conducted specifically for the purpose of this study. Considering the breeding cycle of the sheep and overlaps between generations, our study, conducted in 2022, is composed of lambs after 2 (19%), 3 (53%), 4 (22%) or 5 (6%) generations. Heritability of sociability and human reactivity traits have already been quantified and are available in [33], including the behavioural profiles of high- and low-sociability sheep, which are the main two groups of interest in this study. Overall, highly sociable lambs produce significantly more calls in response to social separation, have a higher locomotion and maintain closer proximity towards conspecifics than less sociable lambs [33].

Lambs used in this study were born outdoors in the spring (from April 2^nd^ to May 2^nd^ 2022), and remained outdoors with their dams until weaning (late June) with limited contact with humans. Lambs had unlimited access to water and pasture and had access to additional food concentrate from four weeks of age prior to selections ran at weaning. Upon selection, selected females lamb had unlimited access to water and pasture and were supplemented with hay (due to the dry 2022 season). Lambs were weighed on June 14^th^ 2022 (average weight: 18.4 ±0.4 kg) and weaned at 75.5 ±8 days of age. Only female lambs, subjected to selection, were considered for this study. From July 4^th^ to 14^th^ 2022, 174 female lambs were subjected to individual tests (average age during the tests: 87 ±9 days), upon which 42 female lambs were selected according to their high (S+, 20 lambs and 1 replacement lamb) or low (S-, 20 lambs and 1 replacement lamb) reactivity to social separation to constitute the two divergent lines of interest (see next section for selection procedure).

Upon selection, the two main groups of S+ and S-lambs were divided into four subgroups of five female within each line (eight groups in total), splitting twins (three times two individuals), mixing individuals of various weights (similar variability in weight at weaning: mean (±sd) of within group standard deviation: 4.31 ±1.3 kg), and avoiding groups fully composed of lambs produced by the same father. Sheep were individually marked within each of their subgroups with color spray and brought to pasture fields (approx. 50*50m). Three lambs (3S+) died from respiratory disease early after the constitution of the groups, one of which was replaced using a replacement lamb, leading to groups of three to five lambs. Eighteen (four groups of three to five) and 20 (four groups of five) lambs, respectively, from the S+ and Sgroups, remained in the study (one lamb selected as replacement for the S-line was not introduced to the groups).

### 2.3 Divergent selection based on behavioural reactivity to isolation and human presence

#### 2.3.1 Tests procedure and behavioural monitoring

The tests were performed on 174 female lambs at the age of 87±9 days. For each test day, the day before being tested, 19-21 female lambs were separated from the rest of the flock of weaned lambs, moved to a barn with both indoor (approximately 15m×8m) and outdoor space (approximately 20m×7 m), and with ad-libitum access to water, hay and concentrate food. The following day, lambs were individually tested in an arena test (morning) and a corridor test (afternoon) in the barn. At the end of each testing day, lambs returned to pasture with the rest of the flock (see below for a summary of the tests; for a detailed description see [39,40]).

##### Arena test

The test arena was a wooden pen (7m long and 2m wide), with the entrance for the tested lamb on one side and three flock-mates placed at the opposite side (contained in a small enclosure), separated by a grid and an optional wooden panel. The test pen was virtually divided into seven zones. In a nutshell, to summarise the procedure used in our study population since 2011, each lamb was gently conducted using wooden panels and barriers by an operator (i.e. without direct contact) in the corridor and was exposed to three consecutive phases: visual contact with the flock-mates (30s), no visual contact with the flock-mates (isolation phase, 60s), and visual contact with the flock-mates while a non-familiar human stood between the tested lamb and the flock-mates (reunion phase, 60s). Several behaviours were measured during both isolation and reunion phases. The number of low and high bleats were recorded manually using an electronic device. Locations were recorded using an automated tracking system allowing to measure the total number of virtual zones crossed and proximity toward conspecifics [41]. The test was also video recorded to score additional behaviours using the Observer 6.0 software (Noldus) by one experienced observer: occurrence of and time spent in vigilance posture, looking at conspecifics, looking at or smelling environment.

##### Corridor test

The test corridor was a non-opaque squared corridor (1m large and 25m long). Each lamb was conducted there by an operator and exposed to two phases: isolation (30s) and exposure to a non familiar moving human (60s). The calls were recorded in this test in the present study but their recordings were not used due to their lower quality, only the occurrence of high bleats in the first phase of the test was used for the selection process (see below). A flight distance between the animal and the moving human was computed in the second phase for genetic selection of divergent lines for reactivity toward humans.

#### 2.3.2 Selection indexes and divergent lines

To summarise the selection process used since 2011, the selection for social traits is based on the reactivity to social separation from conspecifics, and the main variable used in the selection is the number of bleats produced by the lamb during the isolation and reunion phases of the arena test and during the isolation of the corridor test (see [33] for a detailed description). A Principal Component Analysis (PCA) was run on three variables corresponding to the number of vocalisations emitted during each of the three phases of tests to construct a synthetic variable (i.e. first principal component). These three variables all had high loadings on the first component, and accounted equally in the synthetic variable used for genetic selection. For genetic selection of the divergent lines, the Social Selection Index (hereafter ‘SSI’), corresponding to individual estimated breeding values, was calculated using a genetic animal mixed model and considering this synthetic variable and pedigree information [38]. Extremes individuals were chosen based on the SSI to constitute high and low divergent lines. The 42 most extremes individuals for the SSI were selected: the 21 individuals with the highest SSIs, composing the S+ line, *i.e* individuals highly reactive to separation from conspecifics, and the 21 individuals with the lowest SSIs, composing the S-line, *i.e* individuals with low behavioural expression during separation from conspecifics. Intermediate individuals were not selected.

Similarly, two additional divergent lines were selected for human reactivity (see detailed procedure in [33]). A symmetrical pattern was used to constitute the H+ and H-lines by selecting individuals with the highest and lowest Human Selection Indexes (hereafter ‘HSI’), respectively expressing the highest and lowest proximity to the human. The S+ and S-lines were selected for having simultaneously intermediate values for the HSI.

### 2.4 Recording contexts and analysis of vocalisations structure

#### 2.4.1 Isolation bleats

The isolation bleats were recorded in the barn, in which the behavioural tests for selection were carried out. An omnidirectional microphone (Sennheiser MK20), connected to a recorder (Marantz PMD670, frequency sampling 44100 Hz) was placed on the top of the arena at 2.0m from the ground (between 1.5 and 4.5m from the lamb) to record calls produced during the isolation phase of the arena test, when the lamb was fully separated from the flock mates. The context of production of these calls is thus a negative situation of relatively high arousal, with regards to the gregarious behaviour of sheep. In total, 174 females were recorded for their bleats during the behavioural tests described above for genetic selection.

#### 2.4.2 Pre-food treat bleats

Pre-food treat calls were recorded while the lambs remained in their experimental pasture fields outdoors, in the eight subgroups of three to five selected females composed after selection. Each of the eight groups composed of lambs of each line (18S+ and 20S-lambs) were given a barley treat (800g per group) right after shaking a metal barley box, every day between 7:00 and 10:00, at random times and for five weeks. Within a few days, this resulted in an association between the sound and the food treat. Recordings started after one week of habituation to the experimental fields. Every day of the four following weeks, vocalisations produced before the delivery of the barley treat were recorded by the experimenter at group level using an omnidirectional microphone (Sennheiser MK20), connected to a recorder (Marantz PMD670), while located at 3 to 15 meters from the lamb. The identity of the caller was assigned when possible by the experimenter. The context of production of these calls is thus a food anticipation context, which we assume as positive and lower in arousal than the social isolation [42,43]. However, the valence of this context was not validated using emotion indicators.

#### 2.4.3 Acoustic analysis of high bleats

Only high bleats, which were consistently produced in both contexts and by the two divergent lines, were analysed in the present study (hereafter ‘bleats’). Isolation and pre-food treat bleats were analysed separately using the same methodology. All clean calls (not overlapped by any over noise) were kept for acoustic analysis. For the isolation context, from the 174 females tested for their behavioural reaction, 152 produced bleats that could be analysed, which led to 1287 bleats; 322 bleats from 21 females selected as S+, 67 bleats from 14 females selected as S-, and 898 bleats from the remaining 117 females that were recorded but not selected as S+ or S-due to their intermediate SSI (hereafter ‘non-selected’). Regarding the pre-food treat context, among the 37 selected females, 26 produced bleats that could be analysed, which led to 519 bleats; 262 bleats from 14 S+ females (including 64 from unidentified callers within the group), and 257 bleats from 12 S-females (including 52 from unidentified callers within the group).

The process for extracting acoustic parameters was automatically computed using the *extract_chunk_features_dir()* function from *SoundChunk* R-package [44], which compiles several low level functions of the *Seewave* and *tuneR* R package [45–47] (details on computation are available in the electronic supplementary material, section ‘Acoustics’). Fifteen acoustic parameters were extracted to describe the spectro-temporal features of the bleats: duration, energy quartiles (Q25, Q50, Q75 and IQR), spectral flatness, dominant frequency over the duration of the call (mean and standard deviation), fundamental frequency over the duration of the call (mean and standard deviation), mean and relative amplitude of the three first formants (table S1).

It has to be noted that the environment of the two recording contexts (indoor vs. outdoor), and the age difference of the lambs between these two contexts, prevented us from running direct comparison of call structure between contexts.

### 2.5 Statistics

All statistics were performed in R version 4.4.0. [34].

#### 2.5.1 Acoustic differences between selected lines

##### Discriminant acoustic parameters between lines and acoustic scores (LDs)

Suitability of the acoustic dataset, composed of the 15 parameters, for dimension reduction was assessed using a Bartlett test (cortest.bartlett() function, *psych* R package, X_105_=1497, p < 0.001 and X_105_=1707, p < 0.001, respectively, for the isolation and the pre-food treat dataset). Two separate Linear Discriminant Analyses (LDA) were then computed, on the acoustic parameters of calls recorded in each context, using bleats produced by selected S+ and S-sheep, with the line as factor (*lda()* function from *MASS* R package [48]). Indeed, since LDA computations maximize differences between tested groups, this procedure allowed us to extract key potentially discriminant acoustic parameters between the bleats produced by the two lines. Because a warning for collinearity was raised between the two linear discriminant functions produced by the LDAs, only the first LD was kept as a unique composite acoustic score describing the structure of a bleat as a linear combination of the 15 parameters separately for each context (table 1). Therefore, one Acoustic Score (LD) was computed for the isolation calls (isoLD), and one for the pre-food treat calls (foodLD).

**Table 1:** Loadings of acoustic parameters on the first linear discriminant function (LD1) extracted from the linear discriminant function analysis (LDA) performed on isolation calls, leading to the acoustic score ‘isoLD’, and on LD1 extracted from the LDA performed on pre-food treat calls, leading to the acoustic score ‘foodLD’. In bold, loading values having absolute relative loading above 0.4, considered as important descriptors of the linear discriminant function. Underligned acoustic parameters illustrate consistently high loadings across both contexts.

| Acoustic parameter | Loading on isoLD | Loading on foodLD |
| --- | --- | --- |
| duration | **0.515** | -0.174 |
| <u>Q50</u> | **-0.745** | **-0.640** |
| Q25 | 0.131 | **-0.526** |
| Q75 | 0.161 | **0.895** |
| IQR | 0.112 | **-1.000** |
| sfm | -0.172 | **0.932** |
| dfreq_mean | **1.000** | -0.036 |
| dfreq_sd | **-0.866** | -0.051 |
| <u>fo_mean</u> | **-0.895** | **0.488** |
| fo_sd | -0.260 | **-0.545** |
| F1_mean | -0.087 | -0.316 |
| <u>F2_mean</u> | **-0.952** | **0.656** |
| F3_mean | 0.129 | -0.132 |
| <u>F2_amp</u> | **0.926** | **0.639** |
| F3_amp | **-0.430** | -0.175 |

NB: dfreq\_mean and dfreq\_sd for pre-food treat LDA were square-root transformed to approximate normal distribution

##### Predicted acoustic score of isolation bleats produced by non-selected females: test for intermediate bleat structure

Non-selected females, i.e. females that showed intermediate behavioural reaction to the test (intermediate SSI and thus not belonging to extreme individuals selected as S+ or S-), were used as a sub-population to test whether they were also intermediate in terms of the acoustic structure of their isolation bleats. To do so, the output coefficients of the LDA ran on the isolation calls from S+ and Slambs were used to predict the Acoustic Score of the calls produced by non-selected females (*predict()* function from *MASS* R package), a method previously used to study the relative position of an extra group (here the non-selected group) compared to two other ones (here the S- and S+ groups) [49,50]. A LMM (*lmer()* function, *lme4* R package [51]) was then built using the isoLD of all females (S+, S- and non-selected) as a response variable, in order to compare the values of isoLD between selected and non-selected females. In this model, the statistical unit is thus a bleat. Two interacting fixed effects were used: ‘group_line’, a categorical factor with three levels (S+ and S-group of females and the group of non-selected females) in interaction with the weight of the lamb at weaning (a z-scored covariate to increase interpretability [52], hereafter ‘Zweight’). The identity of the genetic father and identity of the lamb emitting the call were used as a nested random factor to control for repeated calls from the same individual. It has to be noted that attempting to add a similar random effect to control for the mother identity led to convergence issues, probably due to the low number twin lambs in the acoustic dataset (only 23% of mothers had more than one lamb). The final model was therefore:

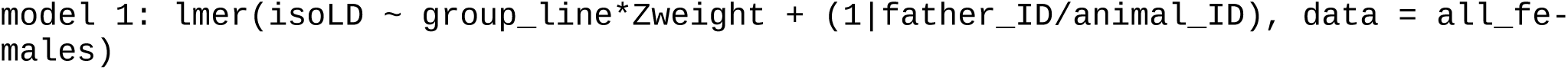

The parallel analysis on pre-food treat bleats could not be performed because non-selected females were excluded from our study upon selection tests, so no pre-food treat calls were recorded for these females.

##### Permuted Discriminant Function Analysis

Permuted Discriminant Function Analyses (pDFA) were used to test if the genetic line of the sheep could be correctly classified above chance level based on the acoustic structure of bleats, while controlling for an unequal number of calls among the two lines within each context and repeated measures. To do so, we focused only on selected sheep, in order to be able to run a parallel analysis in each context of recording (isolation and pre-food treat), using the same 15 acoustic parameters and the line as factor for classification. The statistical approach is described in [53], and the code used here was developed in R by Roger Mundry using the R *MASS* package (built mainly using functions *lda()* and *predict()*), using 100 random selection and cross classification, 1000 permutations to generate p-values for general correct classifications and confusion matrices. Two nested pDFAs per context were ran: the first pDFA (pDFA1) tested for correct classification of all calls without taking into account the individual identity of the emitter (all calls were kept for analysis, even from unidentified callers in the pre-food treat context and from individuals that had produced less than four calls in isolation). The second pDFA (pDFA2) controlled for the individual identity of the lambs: only females of known identity that had produced at least four calls in each context were kept to allow selection of three calls (training dataset) and cross classification on the remaining one(s). For isolation calls, both pDFA included calls from 21 S+ and 14 S-females, with a median number of calls per female [first;last decile] = 9[2;16]. For pre-food treat calls, the first pDFA included calls from 14 S+ and 12 S-females, among which 116 calls out of 519 were produced by unidentified callers, but known line. The second pDFA included calls from seven S+ and nine S-females that had produced at least four calls, with a median number of calls per female [first;last decile] = 12[4;34].

#### 2.5.2 Drivers of acoustic structure of isolation bleats

According to our two hypotheses, isolation bleats structure may be driven by 1) the arousal level expressed during the recording (emotional hypothesis) and/or 2) the selection for sociability (socio-acoustic co-selection hypothesis). The aim of this analysis was to test for potential relationships between isoLD, as a descriptor of bleat acoustic structure, and several potential drivers (selection indexes SSI and HSI, social behaviour, emotional arousal, biometry of the lamb). Because the Acoustic Scores (isoLD) for each context were built to discriminate bleats from S+ and S-females using the bleats of these selected individuals, they were here excluded from this analysis in order to avoid circular reasoning. However, following the previous validation that predicted isoLD of bleats produced by non-selected females were intermediate between S+ and S-ones (see Results), this sub-population of animals could be used instead.

To disentangle between our hypotheses, proxies of arousal and social behaviour were extracted from the scoring of the videos of the isolation and reunion phases to be tested along with the SSI. In a nutshell, behaviours that could be scored in both phases were used to describe the behaviour of S+ and S-lambs: number of virtual zones crossed, occurrence and time looking toward conspecifics (or conspecific door), occurrence and time looking at and smelling environment, and proximity to conspecifics. The computation of a Principal Component Analysis allowed to extract behavioural scores (PCs) that described the behaviour of all tested lamb during isolation (isoPCs) on one side, and during reunion (reuPCs) on the other side (PCA loadings table S2). Then using only S+ and S-lambs in a LMM testing for the effect of the factor line (Type II Anova on LMM), three (out of seven) behavioural scores showed a significant difference or a trend to differ between S+ and S-lamb and were kept as continuous variables potentially explaining bleats structure to be used in the model: IsoPC1 was a proxy of vigilance and mobility (i.e. bodily activation, arousal), ReuPC1 was a proxy of interest toward conspecifics and ReuPC3 was a proxy of attraction toward conspecifics (see loadings of behaviours on the PCs in table S2). More details on the procedure is available in the electronic supplementary material (‘Behavioural profiles’, table S2 and figure S1).

Several controls were added to the model: first we (partially) controlled for potential heritability of the vocal apparatus through the father, using the identity of the father as random factor in the LMM. Indeed, it is likely that the anatomy of the larynx and vocal tract, as well as the way it is operated, is transferred from one generation to the other (leading to similar calls between half siblings), and the identity of the father was the only parameter available to partially control for that. Second, based on the source-filter theory of sound production, larger lambs are expected to produce calls containing lower frequencies [54], which could be a strong predictor of bleat acoustic structure. Hence, the weight of the lamb at weaning (z-scored) was used as interacting covariate in our LMM. Third, to control for effect of the diverging selections, the HSI was also included in our LMM, as a negative control for selections.

Therefore, the full model, using ‘isoLD’ as response variable (and thus bleats as statistical units) was composed of the following fixed effects: isoPC1 (proxy of the arousal of the lamb in isolation, emotional hypothesis), reuPC1 and reuPC3 (proxy of social interest and attraction toward conspecifics, socio-acoustic co-selection hypothesis), the SSI and HSI (proxy of genetic selection for sociability and selection on human reactivity, socio-acoustic co-selection hypothesis), and the z-scored weight of the lamb at weaning Zweight (used as a continuous covariate) in interaction with all other factors. The random factor included the identity of the animal nested within the identity of the father (to partially control for potential heritability of vocal apparatus as well as repetitions of several calls from the same individual). Variance Infection Factors were computed on this model (vif() function, *car* R package) to verify the absence of collinearity between fixed effects (vif values: [min:max]=[1.1:1.5]).

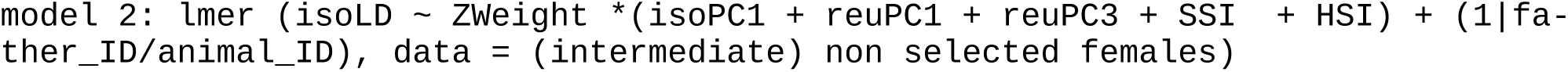

#### 2.5.3 Impact of genetic selection for sociability on encoding of individuality in bleats

To test the impact of the genetic line on identity coding in isolation and pre-food treat bleats, nested pDFAs were computed within each context of recording to extract scores of correct classification according to the identity of the caller.

For the isolation bleats, the calls of 124 females (selected or not) that produced at least four clean calls were used in the analysis. Because, father identity could lead to correct call classifications (table S3A, figure S2A), permutations for testing call classification according to the caller identity (test factor) were restricted to within father identity (restriction factor) ([min;max] = [1;18] females per father for permutations). For the pre-food treat bleats, the calls of 16 females were used in the analysis (with individual identity as a test factor), and due to the lower sample size ([min;max] = [1;3] females per father), no restriction for permutations within father identity was applied. The results of this analysis confirmed that the set of acoustic parameters used in this study allowed for correct classification according to the caller above chance level, in both contexts (electronic supplementary material, table S2), as previously shown in sheep on a different set of parameters [29,55]. The ratio of correctly cross-classified bleats, in each context, was extracted from the diagonal of the prediction/reference confusion matrix as a proxy of success of cross classification of calls according to the identity of the caller (the higher the ratio, the higher the individuality), and used as response variable after square-root transformation in a LMM, in order to test the effect of the genetic line (and potential predictors cited above when possible). The statistical unit is thus the individual.

The LMM built for the isolation bleats included the same potential predictors as above (2.5.2) to explain the correct classification score: SSI, isoPC1, reuPC1, reuPC3 and the weight of the lamb as interacting z-scored covariate (‘Zweight’). Arousal level may decrease identity coding in vocalisations [34]: if identity coding was affected by the level of arousal expressed during the test, then we expected a negative relationship between the ratio of correctly classified calls and isoPC1 (emotional hypothesis). If identity coding was affected by the selection for sociability and social behaviour, then we expected a relationship between the ratio of correctly classified calls and reuPC1, reuPC3 and/or the SSI (socio-acoustic co-selection hypothesis). As in previous models, the father identity was included as a random effect:

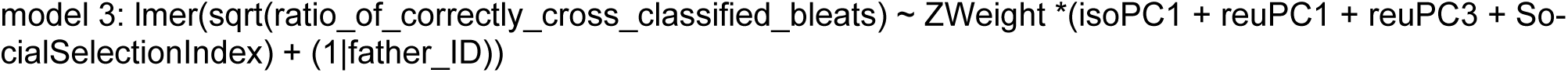

In the LMM built for the pre-food treat bleats, only selected females were included (as non-selected females had not been recorded in this context). Here, and unlike in the model 2 described above, the SSI thus showed a bimodal distribution (extreme SSI selected) and we therefore included the line as a categorical factor (S+ vs. S-) in interaction with the weight of the lamb as an interacting z-scored covariate (‘Zweight’), along with father identity as a random effect. No behavioural data was available in this context to complete the model:

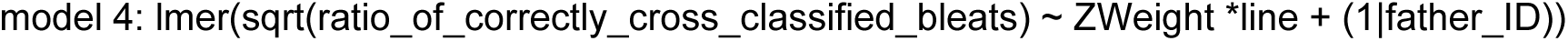

For both models, the response variables were transformed using a square root function to reach symetrical distribution prior to modeling.

#### 2.5.4. Model inspection and statistical tests on LMM

After visual inspection of the normal distribution of the residuals and homoscedasticity (using plot(model), qqPlot(residuals(model)) and hist(residuals(model) functions), a type II Wald Chi-Square test on the full model was computed to test the effect of the potential predictors (Anova() function, *car* R package). Slope estimates were extracted with emtrends() function (‘emmeans’ R package, [56]). When interactions between covariates were significant, pairwise comparisons of slopes were computed, fixing one covariate at mean, mean-se, mean+se to test the variation of the slope of the other covariate. Conditional (R2c) and marginal (R2m) coefficients of determination were computed (r.squaredGLMM() function from *MuMIn* R package [57]). All statistical results of fixed effects-marginal coefficient of determination, multiple comparisons and model estimates are available in the electronic supplementary material.

## 3. Results

### 3.1 Differences in acoustic signature of lambs selected for sociability

Two LDAs were computed using line as factor to build a composite score maximizing the difference between the two lines, separately using isolation and pre-food treat bleats. These LDAs resulted in one Linear Discriminant function (LD1) each, which were then used in further analyses to describe bleat structure (‘Acoustic Scores’, respectively ‘isoLD’ and ‘foodLD’). According to the loadings of acoustic parameters on these LD1s, the median frequency (Q50), the mean fundamental frequency (fo_mean), and the mean frequency of the second formant and its relative amplitude (F2_mean and F2_amp), consistently loaded highly on LD1 in bleats recorded in both contexts (table 1 and figure 1A, 1B). Other parameters loaded highly on LD1 in one of the contexts but not the other: duration and mean dominant frequency (dfreq_mean) loaded more strongly on isoLD, corresponding to isolation bleats, whereas the spectral flatness (sfm), the energy distribution (Q25, Q75 and IQR), and the variability of fundamental frequency (fo_sd), loaded more on foodLD, corresponding to pre-food treat bleats (figure 1B).

**Figure 1:**
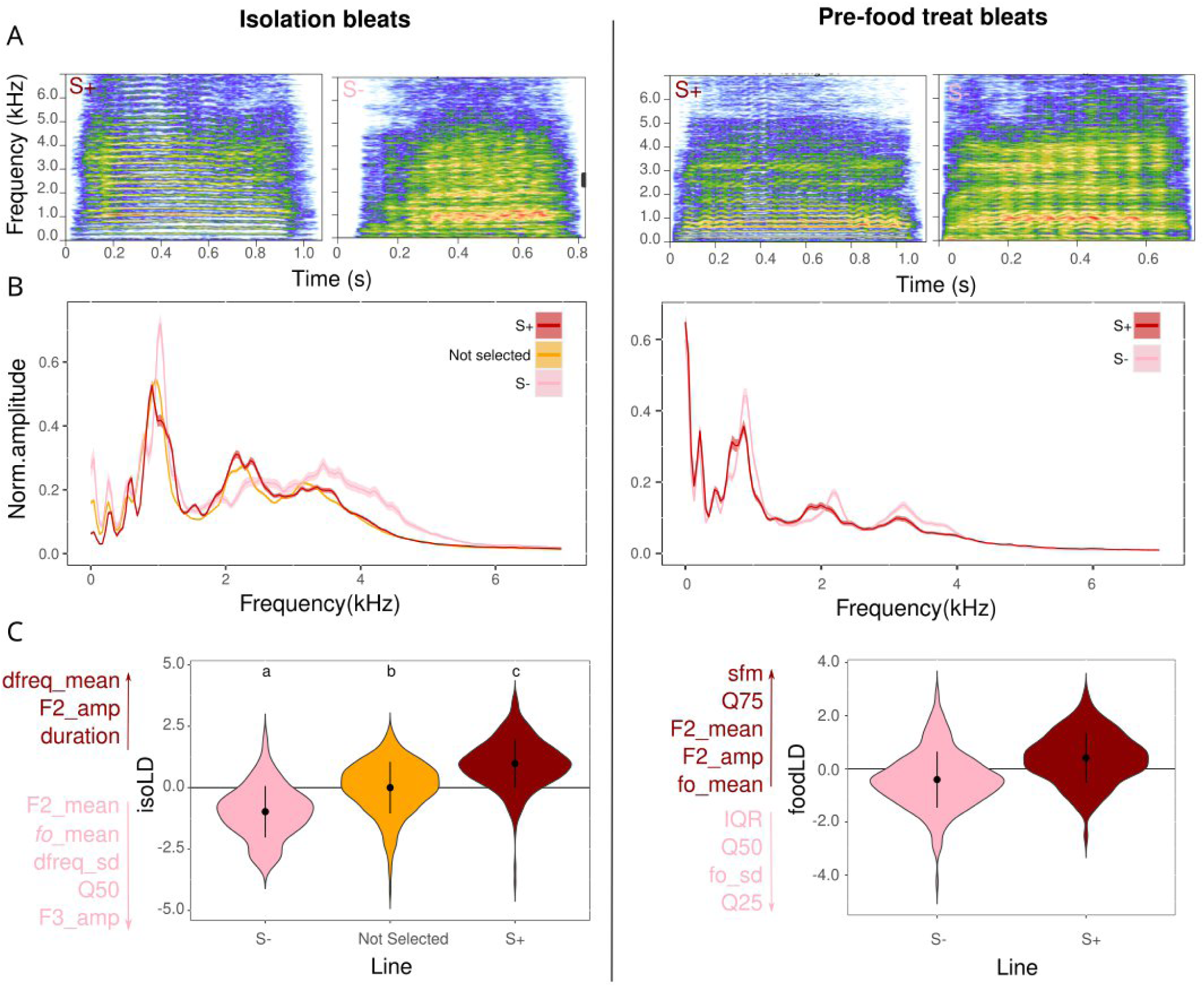
Acoustic differences in isolation bleats (left side of panel) and in pre-food treat bleats (right side of panel) according to the ‘line’ (S+, S- and intermediate not selected when considering the isolation calls). A: Spectrogram of calls of one representative S+ female and one representative S-female according to the context of recording (FFT wl = 2048 pts, ov= 90%, frequency sampling=44.1 kHz, amplitude range from blue to red color scale: -70 to 0dB). B: Averaged mean-spectrum (and standard deviation in shade) over the total amount of calls analyzed per line and contexts (Number of isolation calls: N=322 from S+, N=67 from S-, N=899 from not selected females; Number of pre-food treat calls: N=262 from S+, N=257 from S-females). C: Acoustic parameters explaining the Acoustic Score computed using a LDA on ‘line’ within each context of recording (isoLD for isolation and foodLD for pre-food treat). For isolation calls, the figure also depicts intermediate predictions of isoLD for calls recorded in non-selected females (groups with different letters illustrate statistical differences, see electronic supplementary material, tables S3 for model estimates and pairwise comparisons).

For isolation bleats only, the coefficients of the LDA were used to predict the values of isoLD corresponding to the calls produced by non-selected females and were found having an intermediate position between both extremes S+ and S-individuals (selected on their behaviour) (LMM, Chisq = 23.94, p < 0.001, figure 1C, mean estimate [95%CI]: S-= -0.74[-1.29;-1.19], S+ = 0.7[0.27;1.14], non-selected = -0.09[-0.38;0.21], table S3).

The pDFAs ran separately on isolation and pre-food treat bleats without controlling for individual identity (table 2, pDFA1), showed that the correct classification according to the line was significantly above chance level on the training dataset (24% and 12% correct classification above percentage expected at random for isolation and pre-food treat calls, respectively; p = 0.001 for both; table 2A). However, correct cross classification (i.e. on the validating dataset) was only significantly above chance level for isolation bleats and not pre-food treat bleats (31%, p = 0.001 vs. 15%, p = 0.22; correct cross classification for isolation and pre-food treat calls, respectively; table 2A). Since the results could be driven both by the line and by individual vocal signatures of the lamb belonging to each line, additional pDFAs using individual identity as control factor were performed (pDFA2). For isolation bleats, a trend for correct cross classification above chance level according to the line was found on the validating dataset (p = 0.08), but not for correct classification on the training dataset (p = 0.11). Although the significance 0.05 threshold was not reached for classification in general (table 2A, pDFA2), the confusion matrix of p-values showed significant discrimination of S+ bleats (pDFA2, table 2B). For pre-food treat bleats, no evidence of correct classification of the line was found neither when considering classification scores (p > 0.4, table 2A, pDFA2), nor when inspecting the confusion matrix of p-values for each line (p > 0.3, table 2B, pDFA2).

**Table 2:** Call classification according to the line (S+/S-). pDFAs were run on isolation bleats and on pre-food treat bleats separately with line as factor, and while (pDFA1) or while not (pDFA2) controlling for the individual identity of the caller by included it as a control factor. A: result of pDFAs, percentage of classified and cross-classified calls and p-values computed according to permuted (random) classification. B: confusion matrix of classification p-values for S+ and S-calls in each context.

|  | Isolation bleats |  | Pre-food treat bleats |  |
| --- | --- | --- | --- | --- |
|  | pDFA1 | pDFA2 | pDFA1 | pDFA2 |
| **A. Results of pDFA on discrimination of calls from line S+ and S-** |  |  |  |  |
| Control factor (for training) | X | Individual | X | Individual |
| Number of levels of control factor used for training out of total | X | 14/29 | X | 12/16 |
| Correct classification of training calls (%) | 88.02 | 93.52 | 68.41 | 85.04 |
| Expected correct classification of training calls (chance level) (%) | 64.23 | 85.47 | 56.60 | 84.36 |
| **P value for classification of training calls** | **0.001** | **0.105** | **0.001** | **0.501** |
| Correct cross classification (%) | 80.01 | 68.91 | 64.43 | 56.75 |
| Expected correct cross classification of calls (chance level) (%) | 49.84 | 58.21 | 49.69 | 55.74 |

| P value for cross classification of calls | 0.001 | 0.084 | 0.223 | 0.435 |
| --- | --- | --- | --- | --- |

**B. Confusion matrices of P values of cross classification**
| Reference \ Prediction | S- | S+ | S- | S+ | S- | S+ | S- | S+ |
| --- | --- | --- | --- | --- | --- | --- | --- | --- |
| S- | 0.476 | 0.505 | 0.443 | 0.913 | 0.515 | 0.486 | 0.370 | 0.598 |
| S+ | 1 | **0.001** | 0.947 | **0.039** | 0.875 | 0.126 | 0.586 | 0.459 |

### 3.2 Drivers of acoustic signature of isolation bleats

A significant positive relationship between the SSI and the Acoustic Score isoLD was found (LMM: Chisq = 8.15 p = 0.004, slope estimate [95%CI] = 0.36[0.10;0.63], figure 2), without any interaction with the weight of the lamb (LMM: Chisq = 0.31 p = 0. 576). Based on the acoustic loadings described in table 1, this suggests that non-selected lambs having SSI closer to S+ sheep produced calls with a higher mean dominant frequency, longer duration, and a more pronounced (i.e. of higher relative amplitude) second formant. A significant interaction was found between the weight of the lamb and reuPC3 (LMM: Chisq = 4.29 p= 0.04). Further posthoc tests revealed that there was a positive relationship between reuPC3 and the acoustic score isoLD, especially for heavier lambs (lstrends posthoc tests, slope estimates [95%CI]: 0.27[0.06;0.48] and 0.13[-0.01;0.26] respectively for high and medium), but not for lighter lambs (−0.02[-0.19;0.16], figure 2). According to the loadings of the behavioural PCA (table S2), this suggests that lambs - at least heavy and medium weight ones - exhibiting closer proximity with their conspecifics in the reunion that followed the isolation phase produced bleats with a higher mean dominant frequency, longer duration, and a more pronounced second formant during isolation. There was no evidence for any effect of the behavioural scores isoPC1 - correlated with bodily activation during the isolation - or reuPC1 - correlated with exploration and bodily activation during the reunion - on isoLD (LMM: Chisq < 2.3 p > 0.13, electronic supplementary material, table S6). There was also no evidence of any effect of the HSI on isoLD (LMM: Chisq = 0.006 p = 0.94, electronic supplementary material, table S6).

**Figure 2:**
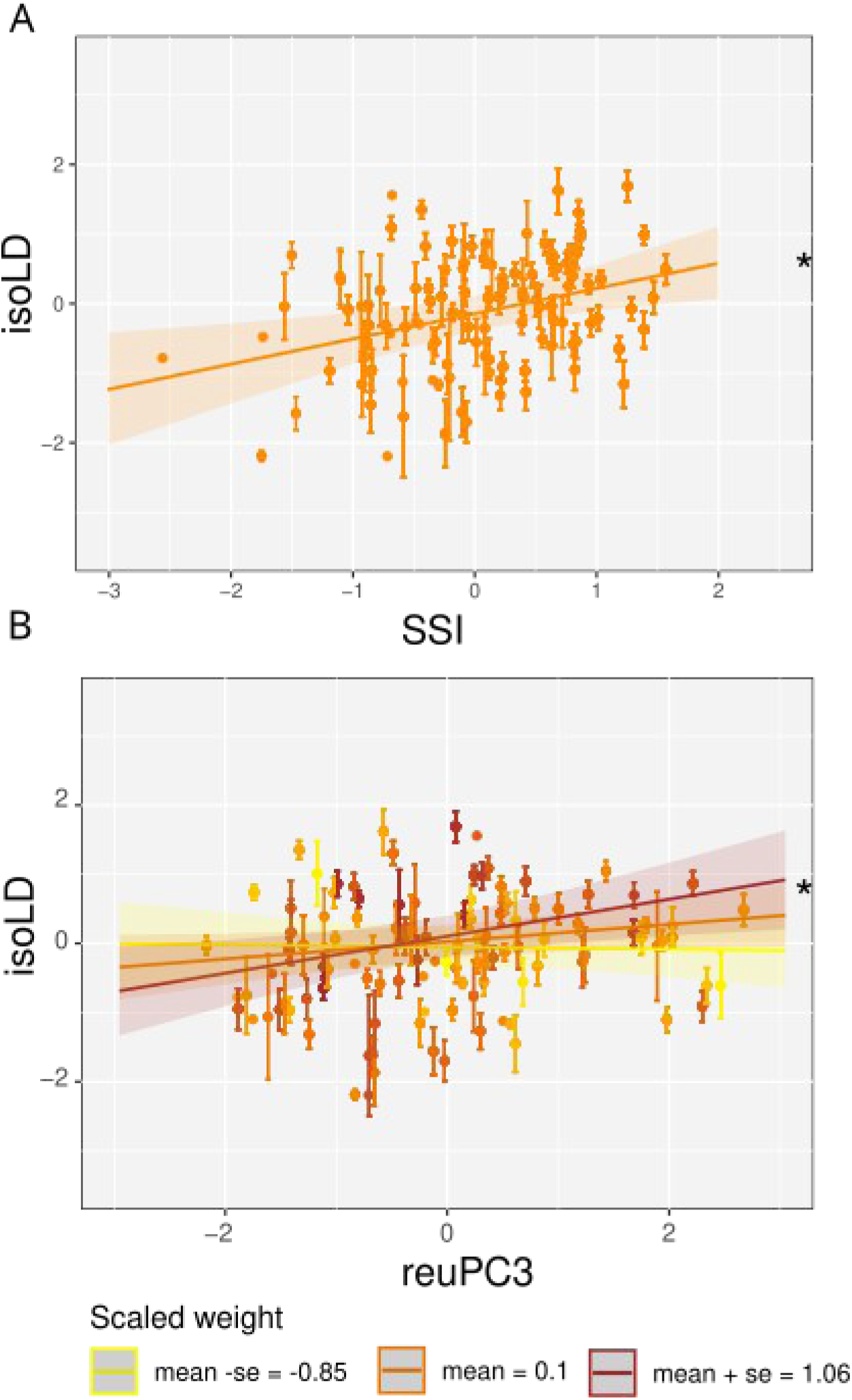
Model estimates and 95% confidence interval depicting the relationship between the Acoustic Score describing isolation bleat structure (isoLD) of non-selected females (hence having intermediate Social Selection Index compared to selected animals) and (A) the Social Selection Index or (B) the behavioural score reuPC3, which positively correlated with proximity. For reuPC3, a significant interaction was found with the weight of the lamb. Therefore, slope estimates are shown based on the mean, mean + se and mean – se of the z-scored weight at weaning used as covariate. Stars (*) show significant slopes. Statistical results of Anova() and model estimates are available in the electronic supplementary material, tables S6 and S7 respectively.

### 3.3 Consequence of genetic divergence for sociability on individuality coding in bleats

For isolation bleats, using 122 females that produced more than four calls during the arena test, the SSI was negatively correlated with the individuality in calls (measured as the ratio of correctly cross-classifed calls in the pDFA) (LMM: Chisq = 4.6 p = 0.03); higher SSI females thus had slightly lower ratios of correctly cross-classified calls (slope estimates [95%CI]: -0.04[-0.09;0.002], figure 3). No evidence of any effect of other potential predictors (Zweight, isoPC1, reuPC1, reuPC3) on call individuality were found (LMM: Chisq < 1.59 p > 0.21, table S8). For pre-food treat bleats, using the 16 females that produced more than four calls during the pre-food treat sessions, no evidence of any effect of the line nor the weight was found on the ratio of correctly cross-classified calls (LMM: Chisq < 0.95 p > 0.33, table S8).

**Figure 3:**
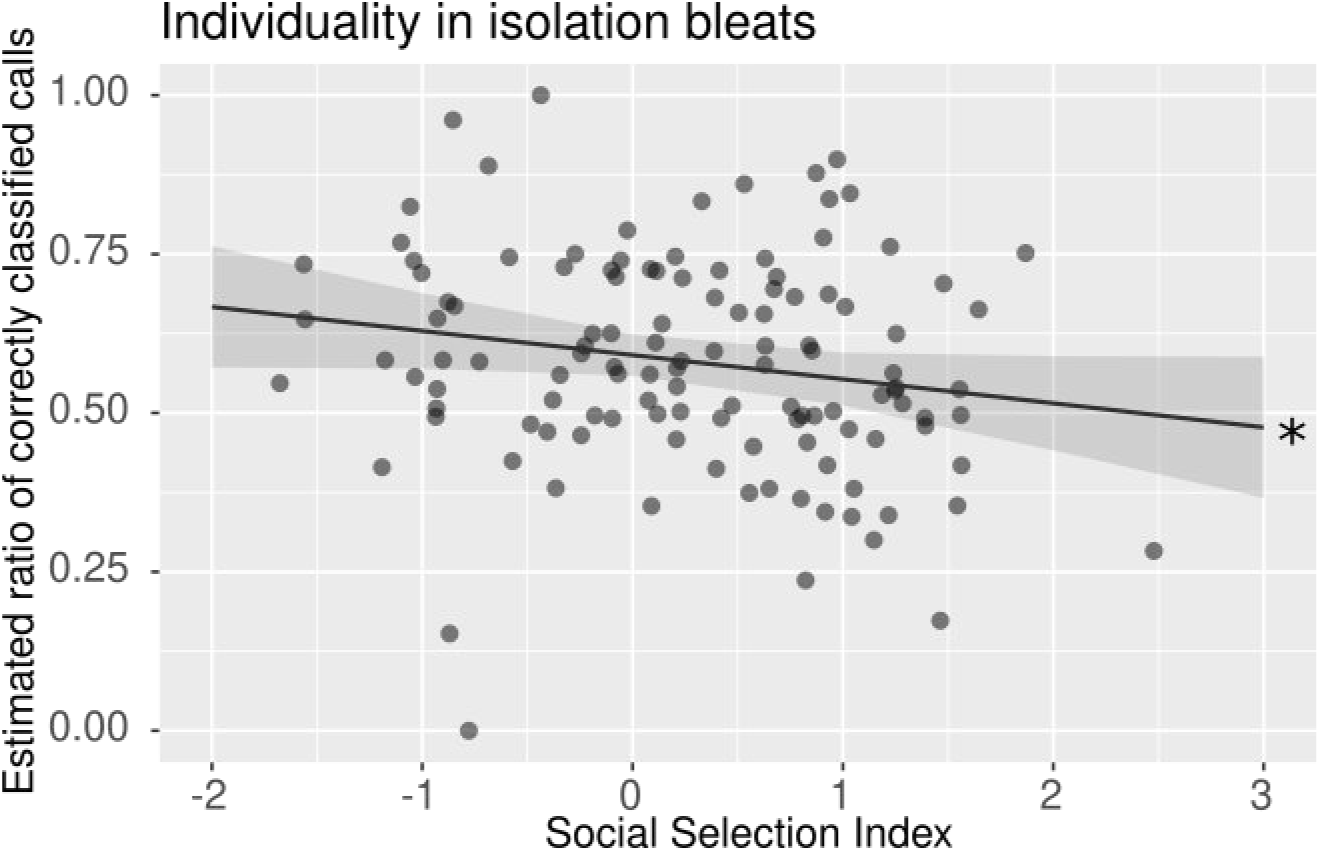
Relationship between the ratio of correctly cross classified calls (proxy of individuality in spectro-temporal features of bleats) and the Social Selection Index. Model estimate (line) and 95% confidence interval represented (shade). Each point represents individual estimated value. *: significant slope. Statistical results of Anova() are available in the electronic supplementary material, tables S8.

## 4. Discussion

High bleats of lambs selected for high (S+) or low (S-) sociability (i.e. reactivity to social isolation) were recorded in emotional contexts, namely isolation and pre-food treat contexts (calls produced at the announcement of a food treat). Our acoustic analysis showed, first, that some acoustic parameters consistently contributed to the difference between the S+ and Slines across contexts, while others were context specific. Second, a line effect on the acoustic structure was particularly present in isolation bleats, while this was not the case in pre-food treat bleats, suggesting a sociability-related ‘line acoustic signature’ that is revealed during the socially challenging situation. Third, the Social Selection Index (SSI) and proximity to conspecifics expressed during the reunion phase significantly impacted the acoustic structure of isolation bleats, while no clear effect of emotional arousal proxies were found. Last, the SSI negatively impacted the encoding of individual’s identity in isolation bleats: highly sociable animals tended to produce isolation bleats with lower probability of being correctly attributed to the caller. Overall, this suggests that temperament traits such as sociability strongly impact on vocalisations of farm animals such as sheep, prevailing in some contexts over the effect of emotional arousal, and impacting on vocal individuality.

### 4.1 Context specific divergence of acoustic structure of bleats in sociable and less sociable sheep lines

The results from the acoustic analysis showed that several acoustic parameters were involved in the discrimination (i.e. loaded highly on isoLD and foodLD) of high bleats produced by female lambs belonging to two divergent genetic lines based on social traits (S+, highly sociable and S-, less sociable) across contexts of recording: mean fundamental frequency, median of the mean spectrum, mean value of second formant and its relative amplitude. Based on the source-filter theory of vocal production, vocal signals result from a two-stage production, with the glottal wave generated in the larynx (the source) being subsequently filtered in the supralaryngeal vocal tract (the filter)[54]. Here, we show that both source (fundamental frequency) and filter (formants) parameters may participate in the discrimination of the divergent lines consistently in both contexts. It is likely that the anatomy of the larynx and the vocal tract and the way it is operated may be influenced by genetic characteristics (heritability of structures of the vocal apparatus), leading to diverging vocal signature between sheep of the two lines (calls from half siblings being more similar, as the correct classification of lamb isolation calls according to the identity of their father showed it, figure S2A, table S3A). However, if genetic was the only mechanism underlying the acoustic difference between our two lines, we would have expected a correct classification of bleats according to the line regardless of the recording contexts. On the contrary, only isolation bleats could be correctly classified according to the line and not pre-food treat bleats. The ‘line acoustic signature’ is hence context-specific, suggesting that genetics of structures of the vocal apparatus cannot, alone, explain the acoustic difference in bleat structure between lines revealed in our study.

This conclusion may find support in the fact that other acoustic parameters participated (i.e. loaded highly on isoLD and foodLD) in the discrimination between the two lines in one context but not the other. Indeed, the dominant frequency mean and its standard deviation, the relative amplitude of the third formant and duration were important parameters for distinguishing S+ from S-only in isolation bleats, but not in pre-food treat bleats. In several species, stress involves the production of calls with longer durations and higher frequencies [3–5]. As sheep are highly gregarious, separation from con-specifics was assumed to be a negative situation of high arousal (provoking high bodily activation [58]). Here, S+ lamb, which expressed a higher behavioural reaction to isolation (see also [33]), also produced high bleats with higher dominant frequency and longer duration. Hence, our acoustic results are in line with the hypothesis that S+ lambs may be more stressed in social isolation than S-lambs. However, specific changes in acoustic structure of bleats as a function of the emotional valence and arousal of the producer are still lacking for sheep, and this should be further validated with standard tests [5].

Different parameters played an important role in distinguishing high bleats from S+ and S-lamb in pre-food treat context: quartiles of the mean spectrum and its inter-quartile range as well as the spectral flatness (noisiness). However, the classification of these calls according to the line was not above chance level. The pre-food treat context may represent an emotional state of intermediate arousal (walking or running), but characterized by a positive valence (pleasant), and not related to a social challenge like in the case of the social separation [42,43]. In fact, it is possible that the selection process, based on sociability, leads to a stronger selective pressure on isolation bleats than on pre-food treat bleats, explaining why the difference emerges in isolation bleats, especially in calls from highly sociable lambs.

### 4.2 Beyond social stress: vocal encoding of heritable sociability in isolation bleats

To investigate whether the above-described ‘line acoustic signature’ is driven by emotional arousal or other factors, we ran a model to test for effects related to the social selection process, the social behaviour expressed during the test and emotional arousal, while controlling for the weight of the animal. Results showed that, although the emotional arousal during isolation was clearly different between the two lines, with S+ being more active (as previously showed on this population using earlier generations [40]), there was no evidence of a relationship between the arousal proxy (i.e. bodily activation) and the acoustic score describing isolation bleats structure. Instead, the Social Selection Index and spatial proximity to conspecifics during the reunion that immediately followed the isolation significantly impacted the acoustic score describing isolation bleat structure. This allowed us to potentially disentangle between the two hypotheses we proposed, aiming at explaining differences in bleat acoustic structure between the two lines of sheep: the emotional hypothesis and the socio-acoustic co-selection hypothesis. Our results provide the first evidence of a vocal encoding of sociability, a temperament trait, which prevails over vocal encoding of emotional arousal in a sheep population selected for sociability. Although more generations and genetic controls would be needed to validate this hypothesis, our result suggest the a ‘socio-acoustic co-selection’: selection for sociability traits could also lead to co-selection of acoustic traits. In other words, selecting for sociability, based on behavioural reaction in response to a social challenge, select for individuals more motivated to be closer to conspecifics after a short term isolation, and may co-select for individuals producing isolation bleats having a typical vocal signature or expressing emotional arousal/motivation to reunite with conspecifics to a higher extent than predicted by their behaviour only. Indeed, first, temperament has been shown to affect expression of emotions in other ungulates such as horses, in which proactive horses appear more stressed based on their behaviour compared to their physiology [8]. Second, in genetic selections, selection for a trait might co-select another one, with several examples in poultry reviewed by Jones and Hocking (1999) [59], a similar mechanism may occur here.

### 4.3 Implications for social cognition

Our results confirmed that an individual vocal signature is encoded in high bleats, in accordance with previous studies [27,29]. Interestingly, based on the acoustic parameters used here, the classification of calls to the correct caller was slightly impacted by the Social Selection Index: the higher the index, and thus the higher the sociability, the lower the ratio of isolation bleats correctly classified to the caller. This result may explain the difference of ‘line acoustic signature’ between the two studied contexts. Indeed, according to the confusion matrix, the ‘line acoustic signature’ was particularly strong in isolation bleats produced by high sociable lambs. Hence, it would make sense that if vocal individuality decreases, calls are more stereotypical, which might lead to a clearer line signature.

Several hypotheses have been proposed to explain individual distinctiveness in calls, summarised in [60]. Based on the ‘distance communication hypothesis’, individually in calls aimed at long-distance communication should be higher than those aimed at short-range communication, for which identity of the caller may be encoded by other modalities (e.g. visual or olfactory [61]). In our study species, high bleats may be considered as rather long-distance calls: contrary to low bleats, they are produced at higher amplitude and when an individual loses contact with conspecifics (however, short separation distance may also trigger the production of high bleats in experimental settings, [30]). In accordance to the distance communication hypothesis, high bleats encode more individuality than low bleats, at least in adult female sheep [28]. Our results revealed a rather high individuality in the high bleats, as shown by high cross-classification scores in general. In addition, higher scores were found in isolation bleats than pre-food treat bleats, which may be in line with the distance communication hypothesis, since during isolation, sheep were at further distance from conspecifics. Secondly, based on the ‘social hypothesis’, individuality in call structure should be driven either by the context or the receiver the call is intended to: calls addressed to a specific individual are predicted to display higher individuality than calls directed to a group or produced in a general context (food, alarm calls, distress) [62–64]. In baby cries for example, individual signatures in pain cries are less reliable than in discomfort cries and it was hypothesized that low individuality in distress calls produced within a social group function to trigger the attention of the entire group rather than to the caretaker only [34]. Recently, using a database of contact calls from seven ungulate species, Osiecka et al. showed that individual distinctiveness in calls was impaired in negative contexts compared to positive ones of similar arousal [35], and hypothesized that calls produced in negative contexts may be addressed to a broader audience than those produced in affiliative contexts. Our results show that, within the same negative context (social isolation) and beyond social stress, encoding of sociability also impairs individuality in high bleats. Hence, calls produced by more sociable animals in a context of social separation, when they are highly motivated to reunite with conspecifics, carry less individuality and may thus be perceived by multiple receivers and increase the probability of triggering a response from a group member, independently from the identity of the caller. If this hypothesis is true, it could have implications for social cognition, as it might impair individual recognition (mother-lamb or social partners) of more sociable sheep based on their high bleats. More research is needed to investigate the consequences of vocal encoding of sociability on communication and its interplay with vocal perception of emotions and social organisation.

### 4.4 Conclusion

To our knowledge, our study provides the first evidence of the consequences of genetic selection for sociability traits on vocal communication, using sheep as a model species. Our findings lead us to suggest a socio-acoustic co-selection: selecting for sociable animals leads to exaggerated vocal expressions of emotional arousal during social isolation and impairs vocal signatures at the group and individual level. The phenomenon we quantified opens the question of the long-term consequences of such selection on social cognition. The divergent lines we describe here also provide a model to study the co-evolution between social traits and social and emotional communication in a gregarious species and explore welfare consequences.

## Supporting information

Electronic supplementary material

## Acknowledgments and funding

The authors would like to thank Sara Parisot and Charlotte Allain for the support of INRAE Experimental Unit of La Fage, Monica Padilla de la Torre for participating to data collection and Eric Delval (UMRH) for scoring the videos of the arena test. This study was funded by a grant from Carlsberg Foundation attributed to Elodie F. Briefer (CF20-0538).

## ***5*** Authors’ contribution

ASV designed the study, collected the acoustic data, ran the analyses and drafted the manuscript, AB designed the behavioural tests, discussed the design of the study and revised the manuscript, GB, CD and SD carried out the behavioural tests, MMM discussed the design of the study and revised the manuscript, PRG collected the acoustic data, ran the analyses and revised the manuscript, DH designed and computed the genetic selections and revised the manuscript, EFB designed the study, revised the manuscript and funded the study.

## Notes

### Competing Interest Statement

The authors have declared no competing interest.

### Summary of Updates

The revised version follows a round of reviews from two anonymous external independent reviewers.

## References

1. Charlton BD, Pisanski K, Raine J, Reby D. 2020 Coding of Static Information in Terrestrial Mammal Vocal Signals. In Coding Strategies in Vertebrate Acoustic Communication (eds T Aubin, N Mathevon), pp. 115–136. Cham: Springer International Publishing. (doi:10.1007/978-3-030-39200-0_5)

2. Bradbury JW, Vehrencamp SL. 2011 Principles of Animal Communication. Second. See http://www.sinauer.com/principles-of-animal-communication.html.

3. Briefer EF. 2020 Coding for ‘Dynamic’ Information: Vocal Expression of Emotional Arousal and Valence in Non-human Animals. In Coding Strategies in Vertebrate Acoustic Communication (eds T Aubin, N Mathevon), pp. 137–162. Cham: Springer International Publishing. (doi:10.1007/978-3-030-39200-0_6)

4. Briefer EF. 2012 Vocal expression of emotions in mammals: mechanisms of production and evidence. J Zool 288, 1–20. (doi:10.1111/j.1469-7998.2012.00920.x)

5. Villain AS, Briefer EF. in production Vocal signals as indicators of emotions. In Assessment of Animal Welfare - a guide to the valid use of indicators of affective states, United Kingdom: John Wiley & Sons Ltd, Oxford,.

6. Hughes DJ, Kratsiotis IK, Niven K, Holman D. 2020 Personality traits and emotion regulation: A targeted review and recommendations. Emotion 20, 63–67. (doi:10.1037/emo0000644)

7. Keltner D. 1996 Chapter 21 - Facial Expressions of Emotion and Personality. In Handbook of Emotion, Adult Development, and Aging (eds C Magai, SH McFadden), pp. 385–401. San Diego: Academic Press. (doi:10.1016/B978-012464995-8/50022-4)

8. Squibb K, Griffin K, Favier R, Ijichi C. 2018 Poker Face: Discrepancies in behaviour and affective states in horses during stressful handling procedures. Applied Animal Behaviour Science 202, 34–38. (doi:10.1016/j.applanim.2018.02.003)

9. Réale D, Reader SM, Sol D, McDougall PT, Dingemanse NJ. 2007 Integrating animal temperament within ecology and evolution. Biological Reviews 82, 291–318. (doi:10.1111/j.1469-185X.2007.00010.x)

10. Carere C, Welink D, Drent PJ, Koolhaas JM, Groothuis TGG. 2001 Effect of social defeat in a territorial bird (Parus major) selected for different coping styles. Physiology & Behavior 73, 427–433. (doi:10.1016/S0031-9384(01)00492-9)

11. Brunelli SA, Hofer MA. 2007 Selective breeding for infant rat separation-induced ultrasonic vocalizations: Developmental precursors of passive and active coping styles. Behavioural Brain Research 182, 193–207. (doi:10.1016/j.bbr.2007.04.014)

12. Mignon-Grasteau S et al. 2005 Genetics of adaptation and domestication in livestock. Livestock Production Science 93, 3–14. (doi:10.1016/j.livprodsci.2004.11.001)

13. Ambruosi S, De Angelis F, Chou J-Y, Goursot C. 2024 Familiar versus unfamiliar: Revealing the complexity of sociability in pigs. Applied Animal Behaviour Science 275, 106248. (doi:10.1016/j.applanim.2024.106248)

14. Miranda-de la Lama GC, Pascual-Alonso M, Aguayo-Ulloa L, Sepúlveda WS, Villarroel M, María GA. 2019 Social personality in sheep: Can social strategies predict individual differences in cognitive abilities, morphology features, and reproductive success? Journal of Veterinary Behavior 31, 82–91. (doi:10.1016/j.jveb.2019.03.005)

15. Miranda-de la Lama GC, Sepúlveda WS, Montaldo HH, María GA, Galindo F. 2011 Social strategies associated with identity profiles in dairy goats. Applied Animal Behaviour Science 134, 48–55. (doi:10.1016/j.applanim.2011.06.004)

16. Forkman B, Boissy A, Meunier-Salaün M-C, Canali E, Jones RB. 2007 A critical review of fear tests used on cattle, pigs, sheep, poultry and horses. Physiology & Behavior 92, 340–374. (doi:10.1016/j.physbeh.2007.03.016)

17. Koolhaas JM, Korte SM, De Boer SF, Van Der Vegt BJ, Van Reenen CG, Hopster H, De Jong IC, Ruis MAW, Blokhuis HJ. 1999 Coping styles in animals: current status in behavior and stress-physiology. Neuroscience & Biobehavioral Reviews 23, 925–935. (doi:10.1016/S0149-7634(99)00026-3)

18. Gartland LA, Firth JA, Laskowski KL, Jeanson R, Ioannou CC. 2022 Sociability as a personality trait in animals: methods, causes and consequences. Biological Reviews 97, 802–816. (doi:10.1111/brv.12823)

19. Šlipogor V, Gunhold-de Oliveira T, Tadić Z, Massen JJM, Bugnyar T. 2016 Consistent inter-individual differences in common marmosets (Callithrix jacchus) in Boldness-Shyness, Stress-Activity, and Exploration-Avoidance. American Journal of Primatology 78, 961–973. (doi:10.1002/ajp.22566)

20. Naguib M, Schmidt R, Sprau P, Roth T, Floercke C, Amrhein V. 2008 The ecology of vocal signaling: male spacing and communication distance of different song traits in nightingales. Behav. Ecol. 19, 1034–1040. (doi:10.1093/beheco/arn065)

21. Carter AJ, Marshall HH, Heinsohn R, Cowlishaw G. 2012 How not to measure boldness: novel object and antipredator responses are not the same in wild baboons. Animal Behaviour 84, 603–609. (doi:10.1016/j.anbehav.2012.06.015)

22. Friel M, Kunc HP, Griffin K, Asher L, Collins LM. 2016 Acoustic signalling reflects personality in a social mammal. Royal Society Open Science 3, 160178. (doi:10.1098/rsos.160178)

23. Nelson XJ, Wilson DR, Evans CS. 2008 Behavioral Syndromes in Stable Social Groups: An Artifact of External Constraints? Ethology 114, 1154–1165. (doi:10.1111/j.1439-0310.2008.01568.x)

24. Leliveld LMC, Düpjan S, Tuchscherer A, Puppe B. 2017 Vocal correlates of emotional reactivity within and across contexts in domestic pigs (Sus scrofa). Physiology & Behavior 181, 117–126. (doi:10.1016/j.physbeh.2017.09.010)

25. Guidi A, Gentili C, Scilingo EP, Vanello N. 2019 Analysis of speech features and personality traits. Biomedical Signal Processing and Control 51, 1–7. (doi:10.1016/j.bspc.2019.01.027)

26. Dwyer CM. 2008 The welfare of sheep. Dordrecht: Springer.

27. Sèbe F, Poindron P, Ligout S, Sèbe O, Aubin T. 2018 Amplitude modulation is a major marker of individual signature in lamb bleats. Bioacoustics 27, 359–375. (doi:10.1080/09524622.2017.1357146)

28. Sèbe F, Duboscq J, Aubin T, Ligout S, Poindron P. 2010 Early vocal recognition of mother by lambs: contribution of low- and high-frequency vocalizations. Animal Behaviour 79, 1055–1066. (doi:10.1016/j.anbehav.2010.01.021)

29. Laliotis GP, Papadaki K, Bizelis I. 2023 Ovine vocal individuality expression by ewes and lambs at a late (40 days) post-partum time point. The Journal of the Acoustical Society of America 153, 751–760. (doi:10.1121/10.0017075)

30. Sèbe F, Nowak R, Poindron P, Aubin T. 2007 Establishment of vocal communication and discrimination between ewes and their lamb in the first two days after parturition. Developmental Psychobiology 49, 375–386. (doi:10.1002/dev.20218)

31. Sebe F, Ligout S, Porter R. 2004 Vocal discrimination of kin and non-kin agemates among lambs. Behaviour 141, 355–369. (doi:10.1163/156853904322981905)

32. Atkinson L, Doyle RE, Woodward A, Jongman EC. 2022 Behavioural reactivity testing in sheep indicates the presence of multiple temperament traits. Behavioural Processes 201, 104711. (doi:10.1016/j.beproc.2022.104711)

33. Hazard D et al. 2022 118. Divergent genetic selections for social attractiveness or tolerance toward humans in sheep. In Proceedings of 12th World Congress on Genetics Applied to Livestock Production (WCGALP), pp. 520–523. Rotterdam, the Netherlands: Wageningen Academic Publishers. (doi:10.3920/978-90-8686-940-4_118)

34. Corvin S, Fauchon C, Patural H, Peyron R, Reby D, Theunissen F, Mathevon N. 2024 Pain cues override identity cues in baby cries. iScience 27. (doi:10.1016/j.isci.2024.110375)

35. Osiecka AN, Lefèvre R, Briefer EF. 2026 Emotional contexts influence vocal individuality in ungulates. Animal Behaviour 231, 123405. (doi:10.1016/j.anbehav.2025.123405)

36. In press. Unité Expérimentale INRAE de la Fage. See 10.15454/1.548325523466425E12 (accessed on 23 April 2026).

37. González-García E, Gozzo de Figuereido V, Foulquie D, Jousserand E, Autran P, Camous S, Tesniere A, Bocquier F, Jouven M. 2014 Circannual body reserve dynamics and metabolic profile changes in Romane ewes grazing on rangelands. Domestic Animal Endocrinology 46, 37–48. (doi:10.1016/j.domaniend.2013.10.002)

38. Hazard D, Bouix J, Chassier M, Delval E, Foulquié D, Fassier T, Bourdillon Y, François D, Boissy A. 2016 Genotype by environment interactions for behavioral reactivity in sheep1. Journal of Animal Science 94, 1459–1471. (doi:10.2527/jas.2015-0277)

39. Boissy A, Bouix J, Orgeur P, Poindron P, Bibé B, Neindre PL. 2005 Genetic analysis of emotional reactivity in sheep: effects of the genotypes of the lambs and of their dams. Genet. Sel. Evol. 37, 381–401. (doi:10.1051/gse:2005007)

40. Ligout S, Foulquié D, Sèbe F, Bouix J, Boissy A. 2011 Assessment of sociability in farm animals: The use of arena test in lambs. Applied Animal Behaviour Science 135, 57–62. (doi:10.1016/j.applanim.2011.09.004)

41. Dore A et al. 2020 A non-invasive radar system for automated behavioural tracking: application to sheep. 2020.12.09.418038. (doi:10.1101/2020.12.09.418038)

42. Briefer EF, Tettamanti F, McElligott AG. 2015 Emotions in goats: mapping physiological, behavioural and vocal profiles. Animal Behaviour 99, 131–143. (doi:10.1016/j.anbehav.2014.11.002)

43. Boissy A et al. 2007 Assessment of positive emotions in animals to improve their welfare. Physiology & Behavior 92, 375–397. (doi:10.1016/j.physbeh.2007.02.003)

44. Villain AS, Renaud-Goud P. 2023 SoundChunk R Package and example data. (doi:10.5281/zenodo.10796326)

45. Sueur J, Aubin T, Simonis C. 2008 Seewave, a free modular too for sound analysis ans synthesis. Bioacoustics 18, 213–226. (doi:10.1080/09524622.2008.9753600)

46. Sueur J. 2018 Sound Analysis and Synthesis with R. Cham: Springer International Publishing. (doi:10.1007/978-3-319-77647-7)

47. Ligges U, Krey S, Mersmann O, Schnackenberg S. 2023 tuneR: Analysis of Music and Speech. See https://CRAN.R-project.org/package=tuneR.

48. Ripley B, Venables B, Bates DM, ca 1998) KH (partial port, ca 1998) AG (partial port, polr) DF (support functions for. 2025 MASS: Support Functions and Datasets for Venables and Ripley’s MASS.

49. Villain AS, Boucaud ICA, Bouchut C, Vignal C. 2015 Parental influence on begging call structure in zebra finches (Taeniopygia guttata): evidence of early vocal plasticity. R Soc Open Sci. 2, 150497. (doi:10.1098/rsos.150497)

50. Villain AS, Hazard A, Danglot M, Guérin C, Boissy A, Tallet C. 2020 Piglets vocally express the anticipation of pseudo-social contexts in their grunts. Sci Rep 10, 18496. (doi:10.1038/s41598-020-75378-x)

51. Bates D, Mächler M, Bolker B, Walker S. 2014 Fitting Linear Mixed-Effects Models using lme4. arXiv:1406.5823 [stat]

52. Schielzeth H. 2010 Simple means to improve the interpretability of regression coefficients. Methods in Ecology and Evolution 1, 103–113. (doi:10.1111/j.2041-210X.2010.00012.x)

53. Mundry R, Sommer C. 2007 Discriminant function analysis with nonindependent data: consequences and an alternative. Animal Behaviour 74, 965–976. (doi:10.1016/j.anbehav.2006.12.028)

54. Taylor AM, Reby D. 2010 The contribution of source–filter theory to mammal vocal communication research. Journal of Zoology 280, 221–236. (doi:10.1111/j.1469-7998.2009.00661.x)

55. Sèbe F, Poindron P, Ligout S, Sèbe O, Aubin T. 2018 Amplitude modulation is a major marker of individual signature in lamb bleats. Bioacoustics 27, 359–375. (doi:10.1080/09524622.2017.1357146)

56. Lenth RV. 2016 Least-Squares Means: The R Package lsmeans. 69, 1–33. (doi:doi:10.18637/jss.v069.i01)

57. Bartoń K. 2016 MuMIn: Multi-Model Inference. R package version 1.15.6.

58. Bradley MM, Codispoti M, Cuthbert BN, Lang PJ. 2001 Emotion and motivation I: defensive and appetitive reactions in picture processing. Emotion 1, 276–298. (doi:10.1037/1528-3542.1.3.276)

59. Jones RB, Hocking PM. 1999 Genetic Selection for Poultry Behaviour: Big Bad Wolf or Friend in Need? Animal Welfare 8, 343–359. (doi:10.1017/S0962728600021977)

60. Linn SN, Schmidt S, Scheumann M. 2021 Individual distinctiveness across call types of the southern white rhinoceros (Ceratotherium simum simum). Journal of Mammalogy 102, 440–456. (doi:10.1093/jmammal/gyab007)

61. Mitani JC, Gros-Louis J, Macedonia JM. 1996 Selection for acoustic individuality within the vocal repertoire of wild chimpanzees. Int J Primatol 17, 569–583. (doi:10.1007/BF02735192)

62. Snowdon CT, Hausberger M. 1997 Social Influences on Vocal Development. Cambridge University Press.

63. Lemasson A, Hausberger M. 2011 Acoustic variability and social significance of calls in female Campbell’s monkeys (Cercopithecus campbelli campbelli). The Journal of the Acoustical Society of America 129, 3341–3352. (doi:10.1121/1.3569704)

64. Papadaki K, Laliotis GP, Bizelis I. 2021 Acoustic variables of high-pitched vocalizations in dairy sheep breeds. Applied Animal Behaviour Science 241, 105398. (doi:10.1016/j.applanim.2021.105398)

