## Supplementary material for "Vocal encoding of heritable sociability prevails over emotions in sheep bleats": Electronic supplementary material

### Additional content

#### *Figure S0: Methodological figure of the study*

*In rows are presented: the procedures applied during the study, in a chronological order, the type of data/variable collected and the number of animals used in each procedure. In columns at the bottom [1:4] illustrate procedures (experimental design or analysis strategy).*

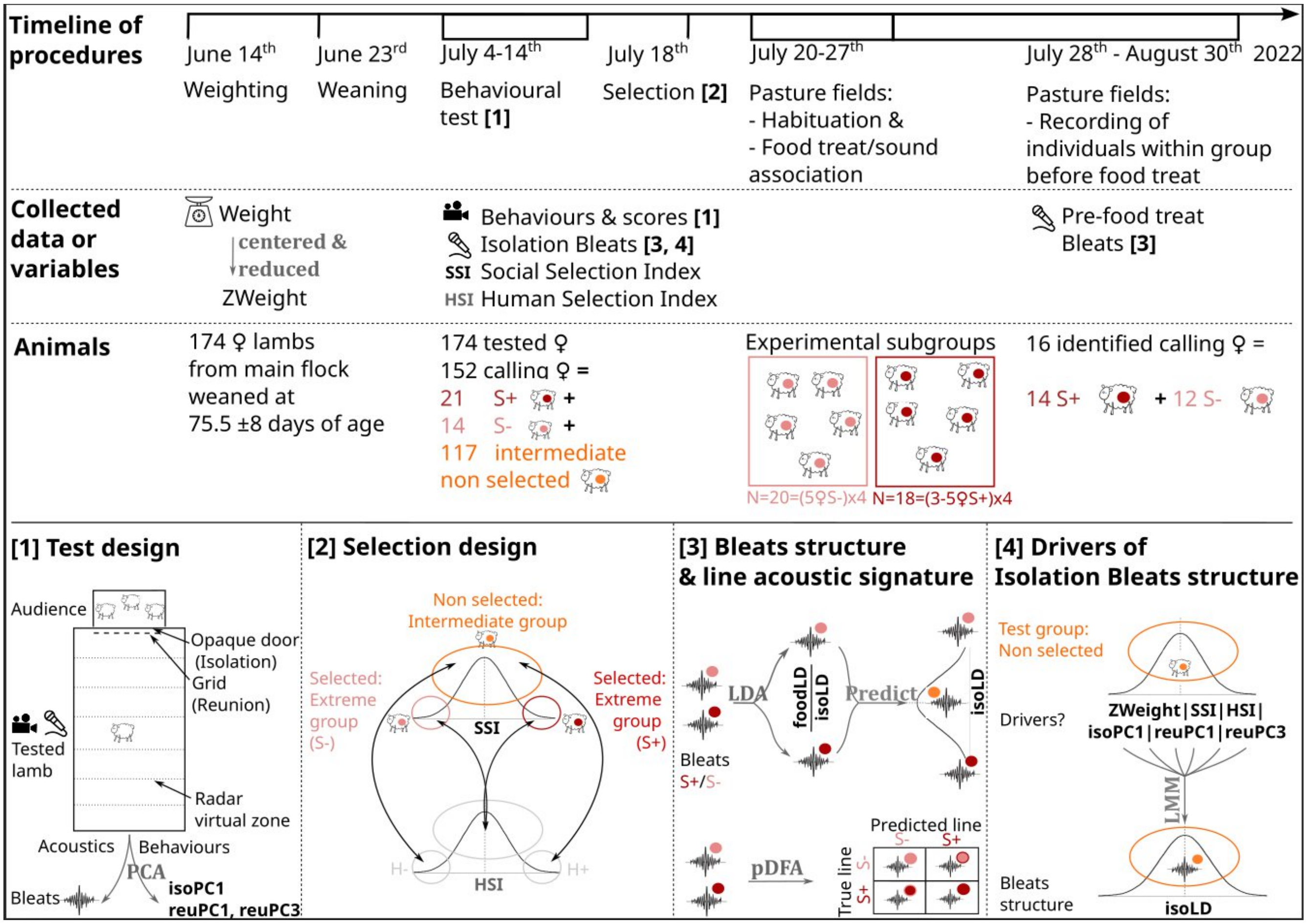

#### *Acoustics*

##### Details on extraction for acoustic parameters

Acoustic parameters were automatically extracted using the *extract_chunk_features_dir()* function of the SoundChunk R-package, based on several low level functions of the Seewave R package [1]. The extraction was computed as follows: the sound was first band-pass filtered between 100-7000 Hz (Fast Fourier Transform FFT hanning window length wl=1024 pts and overlap ov=50%). The duration of the call was then determined upon its detection based on a 5% amplitude threshold on the smoothed en-

velop (wl=4000 pts, ov=90%). After computation of the mean-spectrum (FFT wl=1024pts, ov=50%), parameters related to the general frequency distribution were extracted: first (Q25), second (Q50) and third (Q75) quartiles, Inter-Quartile-Range (IQR) as well as the Wiener entropy (sfm). The dominant frequency was computed on each temporal sliding window (FFT wl=1024 pts and ov=50%), with a 10% amplitude threshold, by searching the loudest frequency in each window. The average of the dominant frequencies (dfreq_mean) and the standard deviation (dfreq_sd) were extracted. The fundamental frequency (*fo*) was computed using the same procedure as the dominant frequency, but after a 100-400 Hz filtering of the calls. The average (*fo*_mean) and standard deviation (*fo*_sd) were extracted. Finally, a formant analysis was performed. First, three frequency peaks were searched after computing a rough mean-spectrum (FFT wl=256 pts, ov=50%), following filtering of the call between 400-7000 Hz. The relative amplitude of each frequency peak was extracted (F1_amp, F2_amp, F3_amp). Based on the values of these three peaks, three frequency bands were used to extract the mean dominant frequency over the duration of the call for each expected formant: the call was successively filtered within each frequency band and the average of the dominant frequencies were extracted (F1_mean, F2_mean, F3_mean). The process was fully automated and hence did not involved visual inspection for decision on duration or position of the formants.

#### Table S1: Acoustic parameters

*Acoustic parameters automatically extracted for spectro-temporal analysis of each bleat. Name, unit, definition and R functions used to extract them is indicated.*

| Acoustic parameter | Unit | Description | Seewave functions |
|---|---|---|---|
| duration | s | Duration of the call. Based on detection of 5% of the smoothed envelop energy (smoothing using wl=4000pts, ov=90%) | ffilter(), timer() |
| Q50 | Hz | Median of the mean spectrum (FFT wl=1024 pts, ov=50%) computed within the frequency range 100-7000 Hz | |
| Q25 | Hz | First quartile of the mean spectrum (FFT wl=1024 pts, ov=50%) within the frequency range 100-7000 Hz | ffilter(), meanspec(), specprop() |
| Q75 | Hz | Third quartile of the mean spectrum (FFT wl=1024 pts, ov=50%) within the frequency range 100-7000 Hz | |
| IQR | Hz | Inter-Quartile-Range, difference between the Q75 and the Q25 computed on the mean spectrum (FFT wl=1024 pts, ov=50%) within the frequency range 100-7000 Hz | |
| sfm | | Spectral Flatness, also called Wiener Entropy as a measure of spectral noise (the higher the noisier) | |
| dfreq_mean | Hz | Average of dominant frequencies measured on each sliding window computed over the duration of the call (FFT wl=1024 and ov=50%) within the frequency bandwidth: 100-7000 Hz) | ffilter(), fpeaks(), dfreq() |
| dfreq_sd | Hz | Standard deviation of dominant frequencies measured on each sliding window computed over the duration of the call (FFT wl=1024 and ov=50%) within the frequency bandwidth: | |

| | | | |
|---|---|---|---|
| | | 100-7000 Hz) | |
| fo_mean | Hz | Average of dominant frequencies measured on each sliding window computed over the duration of the call (FFT wl=1024 and ov=50%) after filtering the call between 100-400 Hz, expected frequency range of fundamental frequency. | |
| fo_sd | Hz | Standard deviation of dominant frequencies measured on each sliding window computed over the duration of the call (FFT wl=1024 and ov=50%) after filtering the call between 100-400 Hz, expected frequency range of fundamental frequency | |
| F1_mean | Hz | Average of dominant frequencies measured on each sliding window computed over the duration of the call (FFT wl=1024 and ov=50%, after filtering the calls within frequency bandwidths of putative first, second, and third formant) | |
| F2_mean | Hz | | |
| F3_mean | Hz | | |
| F1_amp | | Normalized amplitude of the first, second or third frequency peak found after computing a mean spectrum (FFT wl=256, ov=50%, frequency bandwidth: 400-7000 Hz). | ffilter(), fpeaks() |
| F2_amp | | | |
| F3_amp | | | |

### ***Behavioural profiles***

#### Table S2: Computation of behavioural scores (PCs)

*Results of the two Principal Component Analyses (PCA) computed on behaviours occurring during the isolation phase (isoPC1, isoPC2, isoPC3, isoPC4) and during the reunion phase (reuPC1, reuPC2, reuPC3) of the arena test (isolation context). Highest loadings for each behaviour appear in bold. The last row depicts significant differences between lambs from S+ and S- divergent lines * p < 0.05, ~ p < 0.10. The three behavioural scores showing trends or significant differences between selected S+ and S- lamb, based on an Anova test on the LMM model, were kept as potential predictors of the Acoustic Score (isoPc1, reuPC1, reuPC3). All statistics and model estimates are available as supplementary material tables S3 and S4 (figure S1 of supplementary material illustrates differences between lines).*

| | **isoPC1** | **isoPC2** | **isoPC3** | **isoPC4** | **reuPC1** | **reuPC2** | **reuPC3** |
|---|---|---|---|---|---|---|---|
| **Eigenvalue** | 3.374 | 1.451 | 1.310 | 1.044 | 2.877 | 1.964 | 1.264 |
| **Cumulative variance explained (%)** | 42.172 | 60.316 | 76.690 | 89.745 | 35.965 | 60.515 | 76.314 |
| Proximity to conspecifics | 5.122 | 0.542 | 14.626 | **76.709** | -6.195 | 20.978 | **68.760** |
| Time spent looking at conspecifics | **-46.299** | 37.229 | 6.162 | 0.643 | **50.312** | -10.010 | -11.066 |
| Occurrence of looks at conspecifics | -31.179 | **36.290** | 20.567 | -5.618 | **-35.782** | -17.429 | -0.033 |
| Occurrence of vigilance posture | -2.278 | 28.283 | **-62.070** | 0.017 | -5.652 | -79.371 | 3.531 |
| Time spent in vigilance posture | **83.495** | 7.582 | -2.531 | 0.211 | -2.086 | **-60.993** | 3.855 |
| Occurrence of looks at or smells environment. | **-43.726** | 0.112 | -24.786 | 21.233 | **-81.066** | 2.429 | -9.194 |
| Time spent looking at or smelling environment. | **-61.511** | -31.925 | -0.245 | 0.006 | **-64.445** | 4.116 | -23.843 |
| Number of virtual zones crossed | **-63.770** | -3.184 | 0.006 | 0.004 | **-42.184** | -1.073 | 6.110 |
| **S+ S- comparison (see also figure S1)** | **S+ < S- *** | **NS** | **NS** | **NS** | **S+ < S- ~** | **NS** | **S+ > S- *** |

To extract proxies of arousal and social behaviour as predictors of bleat acoustic structure, we used behavioural data extracted from automatic recording (i.e. location changes) and from scoring the videos of the last two phases of the arena test: isolation and subsequent reunion with conspecifics. Only behaviours that could be scored in both phases were used: the number of virtual zones crossed, vigilance posture (occurrence and percent of time), looks toward conspecifics/conspecifics's door (occurrence and percent of time), looking at or smelling the environment (occurrence and percent of time), and a proximity score calculated using a formula giving more weight to the time spent in the zones nearest to the conspecifics' door (see formula in [2]). Two Principal Component Analyses (PCA) were computed to build composite scores of the lambs' behaviour during the isolation phase (isoPCs) and during the reunion phase (reuPCs) (dudi.pca() function from 'ade4' R package). PCs having an Eigenvalue above one were kept, resulting in four behavioural scores describing the isolation phase, and three behavioural scores describing the reunion phase. Loadings were extracted using the inertia.dudi() function from 'ade4' R package [3] for interpretation of PCs (table S2). All scored females, selected or not (N=152), were used to build the behavioural scores, and differences in behaviour were tested on the subset of selected females of each line. This was done using a linear mixed effects model to test the effect of the 'line' (S+ or S-) in interaction with the scaled weight of the lamb, while using the identity of the father as random effect, on behavioural scores (PCs) (model and statistics are detailed in supplementary material, figure S1, tables S3 and S4). Only PCs showing a significant or marginally significant difference between S+ and S- were kept as potential predictors. Accordingly, out of seven PCs, three were kept as behavioural descriptors, one for isolation and two for reunion (S+ vs. S-: $Chisq > 3.702$, $p < 0.055$, table S3). IsoPC1, explaining for 42% of the variance, was mainly positively correlated with vigilance and negatively with locomotion during the isolation phase (i.e. bodily activation, table 2). ReuPC1, explaining 35% of the variance, was positively correlated with the time spent looking at conspecifics and negatively to looking and smelling the environment and locomotion (i.e. interest toward conspecifics vs. exploration). Finally, ReuPC3, explaining 15% of variance, was positively correlated with proximity toward conspecifics (i.e. attraction). The behavioural results show that S+ lambs express a higher bodily activation in isolation, a higher exploration during a reunion with conspecifics and a higher

proximity to conspecific than S- lambs (see supplementary material, figure S1), which is in accordance with previous studies on lambs from previous generations of this selection line [4,5].

#### Figure S1: Comparison of isoPC1, reuPC1, reuPC2

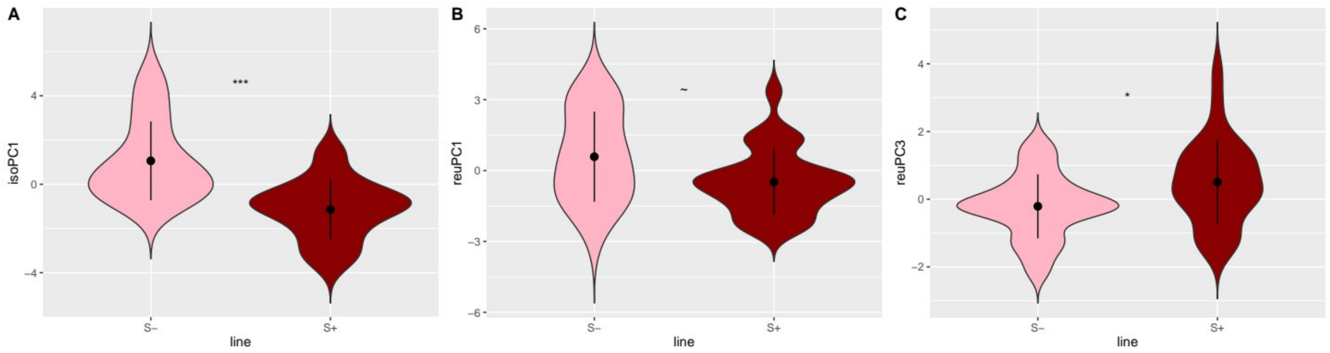

*Violin plots showing significant differences in behavioural scores isoPC1 (A: ***), reuPC1 (B, ~) and reuPC3 (C, *) between S+ and S- during the arena test. No evidence of any effect of the line was found for IsoPC2, isoPC3, isPC4 and reuPC2. Statistical results and model estimates are available in Tables S3 and S4. ***: p< 0.001, *: p< 0.05, ~: p<0.06. Results are consistent with findings on lambs from the same selection process in previous studies [2].*

### ***Individuality***

#### Table S3: Individual classifications

*Call classification according to individuals using pDFA on isolation bleats (A and B, test factor respectively fatherID and callerID) and pre-food treat bleats (B, test factor callerID). Following correct classification of fatherID, restricting permutations to fatherID was applied when possible (B. Isolation calls). Percentage of classified and cross-classified calls and p-values computed according to permuted (random) classification.*

| **A. test factor = father identity (heritability of call structure)** | **Isolation bleats** |
|---|---|
| Restriction factor for permutations | None |
| Correct classification of training calls (%) | 63.015 |
| Expected correct classification of training calls (chance level) (%) | 55.437 |
| **P-value for classification of training calls** | 0.059 |
| Correct cross classification (%) | 23.171 |
| Expected correct cross classification of calls (chance level) (%) | 13.831 |
| **P-value for cross classification of calls** | 0.001 |

| **B. test factor = caller identity (individual vocal signature)** | **Isolation bleats** | **pre-food treat bleats** |
|---|---|---|
| Restriction factor for permutations | Father | None |
| Correct classification of training calls (%) | 80.685 | 92.542 |

| | | |
|---|---|---|
| Expected correct classification of training calls (chance level) (%) | 41.692 | 80.955 |
| **P-value for classification of training calls** | **0.001** | **0.021** |
| Correct cross classification (%) | 35.822 | 22.832 |
| Expected correct cross classification of calls (chance level) (%) | 2.917 | 6.268 |
| **P-value for cross classification of calls** | **0.001** | **0.001** |

#### Figure S2: Confusion matrix of classification of individuals using isolation bleats

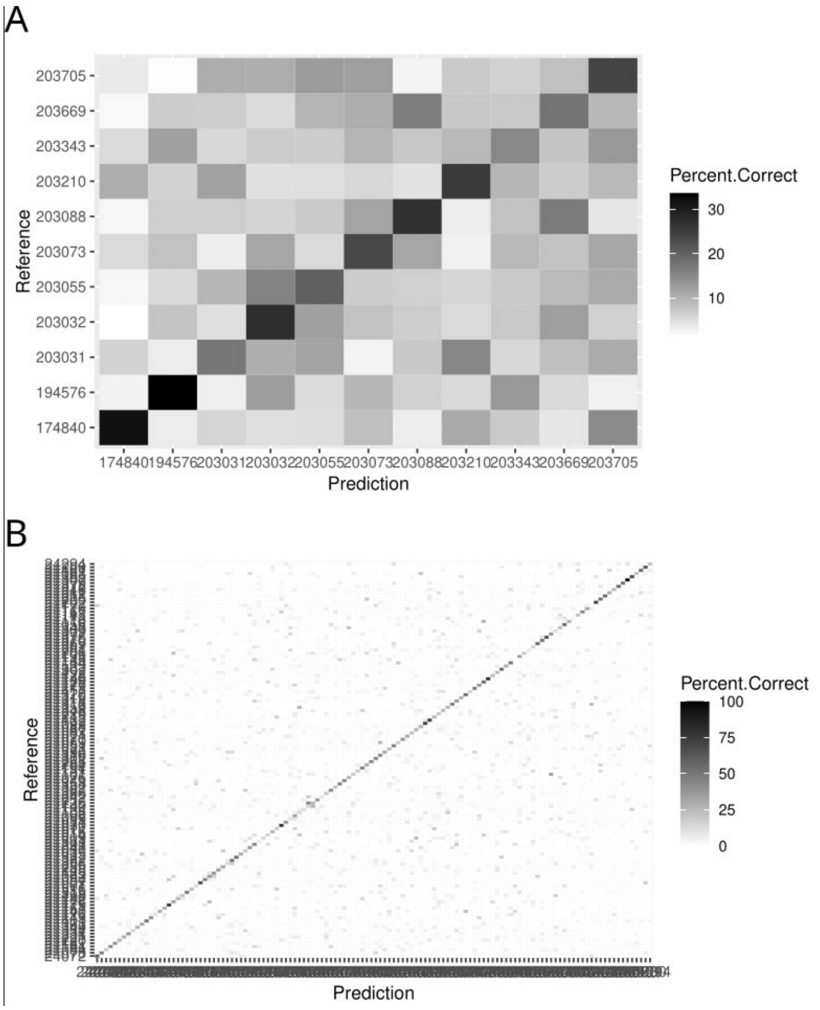

*Reference/Prediction confusion matrix on cross classification of isolation bleats according to the individual (A = father identity, B = caller identity). Real identity of the animal is on the y axis and predicted identity of the analysis using acoustic parameters is on the x axis. Each cell of the matrix is the normalized number of cross classification of a call. The diagonal then represents the correct cross classification of the calls according to the individual: the darker the cell the higher the ratio (form 0 to 100% of cross classification). Following significant classification according to father identity, pDFA for caller identity was computed restricting permutations within father ID.*

#### Figure S3: Confusion matrix of classification of individuals using pre-food treat bleats

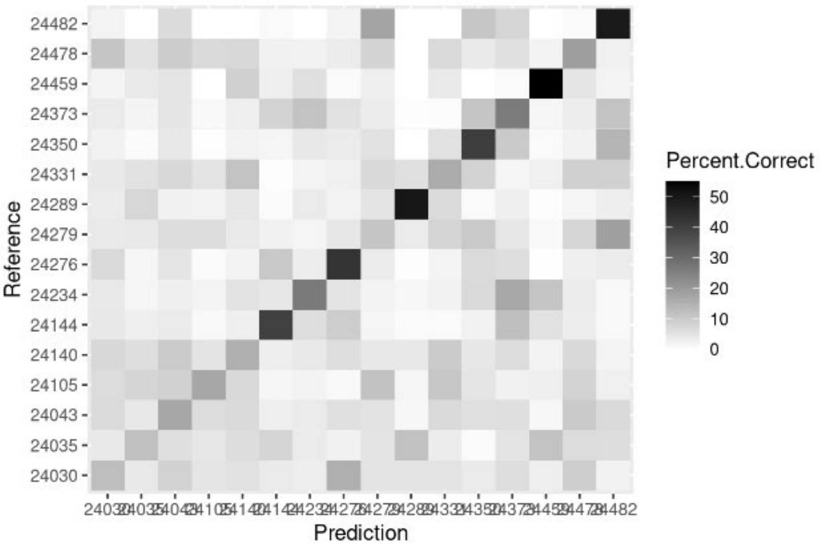

*Reference/Prediction confusion matrix on cross classification of pre-food treat bleats according to the individual. Real identity of the animal is on the y axis and predicted identity of the analysis using acoustic parameters is on the x axis. Each cell of the matrix is the normalized number of cross classification of a call. The diagonal then represents the correct cross classification of the calls according to the individual: the darker the cell the higher the ratio (from 0 to 50% of cross classification).*

### Statistical tables

##### *Table S3: Comparison of Acoustic Score of isolation bleats (isoLD1) between lines.*

*Type II Anova computed on full model with Anova() function ('car' R package). Contrast and model estimates computed with emmeans() function ('emmeans' R package with Tukey contrasts).*

Model 1: lmer(isoLD1 ~ ZWeight* Line + (1 | father_id/animal_id)). *R2c = 0.53, R2m=0.12.*

| **Anova** | **Fixed effect** | **Chisq** | | **Df** | **Pvalue** |
|---|---|---|---|---|---|
| | Zweight | 2.290 | | 1 | 0.130 |
| isoLD1 | Line (3 levels) | 23.938 | | 2 | <0.001 |
| | Zweight:Line | 0.527 | | 2 | 0.768 |
| **Contrast** | **Estimate** | **SE** | **df** | **t.ratio** | **P.value** |
| (1.S-) - (2.Not-Selected) | -0.649 | 0.257 | 173.928 | -2.527 | 0.033 |
| (1.S-) - (3.S+) | -1.437 | 0.326 | 140.300 | -4.407 | <0.001 |

| | | | | | |
|---|---|---|---|---|---|
| (2.Not-Selected) - (3.S+) | -0.788 | 0.208 | 99.335 | -3.797 | 0.001 |

| Group estimates | emmean | SE | df | lower.CL | upper.CL |
|---|---|---|---|---|---|
| 1.S- | -0.735 | 0.273 | 93.089 | -1.278 | -0.192 |
| 2.Not-Selected | -0.086 | 0.134 | 10.661 | -0.381 | 0.209 |
| 3.S+ | 0.702 | 0.214 | 36.040 | 0.269 | 1.135 |

#### *Tables S4 and S5: Comparison of behavioural score between lines*

Linear mixed effect model (lmer() function from 'lme4' R package) testing the effect of line on Behavioural Scores computed using PCAs. The scaled weight was added as interacting fixed effect to control for effect of size (weight and age were correlated) on mobility and social scores. The genetic father (father_id) was added as random factor to account for repeated measures.

```
Model: lmer(PC ~ Line*Zweight +(1|father_ID), data=only selected females)
```

##### Table S4: Anova table

*Type II Anova computed on the full model using Anova() function from 'car' R package.*

| Response variable | R2c | R2m | Fixed effect | Chisq | Df | Pvalue |
|---|---|---|---|---|---|---|
| **isoPC1** | 0.36 | 0.35 | Zweight | 1.005 | 1 | 0.316 |
| | | | **Line** | **24.063** | **1** | **<0.001** |
| | | | Zweight:Line | 0.757 | 1 | 0.384 |
| isoPC2 | 0.01 | 0.04 | Zweight | 0.396 | 1 | 0.529 |
| | | | Line | 0.214 | 1 | 0.644 |
| | | | Zweight:Line | 0.015 | 1 | 0.901 |
| isoPC3 | <0.001 | <0.001 | Zweight | 0.001 | 1 | 0.971 |
| | | | Line | 0.002 | 1 | 0.964 |
| | | | Zweight:Line | 0.001 | 1 | 0.982 |
| isoPC4 | 0.08 | 0.08 | Zweight | 1.092 | 1 | 0.296 |
| | | | Line | 0.726 | 1 | 0.394 |
| | | | Zweight:Line | 2.084 | 1 | 0.149 |
| **reuPC1** | 0.18 | 0.10 | Zweight | 0.022 | 1 | 0.881 |
| | | | **Line** | **3.702** | **1** | **0.054** |
| | | | Zweight:Line | 0.079 | 1 | 0.779 |
| reuPC2 | 0.09 | 0.03 | Zweight | 1.194 | 1 | 0.274 |

| | | | | | | |
|---|---|---|---|---|---|---|
| | | | Line | 0.269 | 1 | 0.604 |
| | | | Zweight:Line | 0.233 | 1 | 0.629 |
| | | | Zweight | 0.005 | 1 | 0.942 |
| **reuPC3** | 0.12 | 0.11 | **Line** | **5.179** | **1** | **0.023** |
| | | | Zweight:Line | 1.240 | 1 | 0.265 |

#### Table S5: Estimates table

*Model estimates computed on the full model (emmeans() function 'emmeans' R package) when a significant effect of 'line' was found.*

| Response.variable | line | emmean | SE | df | lower.CL | upper.CL |
|---|---|---|---|---|---|---|
| isoPC1 | S- | 1.060 | 0.340 | 2.786 | -0.069 | 2.189 |
| | S+ | -1.213 | 0.354 | 4.329 | -2.168 | -0.257 |
| reuPC1 | S- | 0.624 | 0.443 | 3.282 | -0.721 | 1.970 |
| | S+ | -0.523 | 0.432 | 5.003 | -1.633 | 0.587 |
| reuPC3 | S- | -0.180 | 0.229 | 2.670 | -0.963 | 0.603 |
| | S+ | 0.529 | 0.244 | 4.115 | -0.142 | 1.200 |

### *Tables S6 and S7: Predictors of Acoustic Score of isolation bleats (isoLD1) of non-selected females.*

#### Table S6: Anova table.

*Type II Anova computed on full model with Anova() function ('car' R package).*

```
model 2: lmer (isoLD ~ ZWeight *(isoPC1 + reuPC1 + reuPC3 + SSI  + HSI) + (1|father_ID/animal_ID), data = (intermediate) non selected females)
```

| Response variable | R2c | R2m | Fixed effect | Chisq | Df | Pvalue |
|---|---|---|---|---|---|---|
| | | | Zweight | 0.598 | 1 | 0.439 |
| | | | isoPC1 | 0.029 | 1 | 0.865 |
| | | | reuPC1 | 0.375 | 1 | 0.540 |
| IsoLD1 | 0.51 | 0.10 | reuPC3 | 2.262 | 1 | 0.133 |
| | | | **SocialSelectionIndex** | **8.150** | **1** | **0.004** |
| | | | HumanSelectionIndex | 0.006 | 1 | 0.939 |

| | | | | |
|---|---|---|---|---|
| | Zweight:isoPC1 | 0.540 | 1 | 0.462 |
| | Zweight:reuPC1 | 0.092 | 1 | 0.761 |
| | **Zweight:reuPC3** | **4.293** | **1** | **0.038** |
| | Zweight:SocialSelectionIndex | 0.312 | 1 | 0.576 |
| | Zweight:HumanSelectionIndex | 0.046 | 1 | 0.831 |

#### Table S7: Estimates table

Contrast and model estimates computed with emmeans() function (‘emmeans’ R package with Tukey contrasts) on model 2.

| **Response variable** | **Variable** | **Fixed variable** | **Level of fixed variable** | **Slope trend** | **Df** | **Lower.CL** | **Upper.CL** |
|---|---|---|---|---|---|---|---|
| isoLD1 | SocialSelectionIndex | - | - | 0.362 | 53.590 | 0.095 | 0.629 |
| isoLD1 | ReuPC3 | Zweight | -0.85 | -0.016 | 93.430 | -0.188 | 0.155 |
| | | Zweight | 0.1 | 0.125 | 88.360 | -0.008 | 0.258 |
| | | Zweight | 1.06 | 0.268 | 88.346 | 0.059 | 0.476 |

#### ***Tables S8: Anova table, impact of Selection on individuality in calls***

| **Response variable** | **Fixed effect** | **Chisq** | **Df** | **Pvalue** |
|---|---|---|---|---|
| *A. Isolation bleats* | | | | |
| model 3: lmer(sqrt(ratio_of_correctly_cross_classified_bleats) ~ ZWeight *(isoPC1 + reuPC1 + reuPC3 + SocialSelectionIndex) + (1\|father_ID)). *R2c = 0.08, R2m = 0.08* | | | | |
| ratio of correctly cross classified calls (sqrt) | Zweight | 1.589 | 1 | 0.207 |
| | **SocialSelectionIndex** | **4.610** | **1** | **0.032** |
| | isoPC1 | 0.126 | 1 | 0.723 |
| | reuPC1 | 0.041 | 1 | 0.840 |
| | reuPC3 | 0.297 | 1 | 0.586 |
| | Weight : SocialSelectionIndex | 1.697 | 1 | 0.193 |
| | Weight : isoPC1 | 0.302 | 1 | 0.582 |

| | | | | |
|---|---|---|---|---|
| | Weight : reuPC1 | 0.089 | 1 | 0.766 |
| | Weight : reuPC3 | 0.023 | 1 | 0.879 |

*B. pre-food treat bleats*

model 4: lmer(sqrt(ratio_of_correctly_cross_classified_bleats) ~ ZWeight *line + (1|father_ID)). *R2c = 0.38, R2m = 0.11.*

| | | | | |
|---|---|---|---|---|
| ratio of correctly cross classified calls (sqrt) | Zweight | 0.003 | 1 | 0.955 |
| | Line | 0.755 | 1 | 0.385 |
| | Zweight : Line | 0.954 | 1 | 0.329 |

### References

1. Villain AS, Renaud-Goud P. 2023 SoundChunk R Package and example data. (doi:10.5281/zenodo.10796326)

2. Ligout S, Foulquié D, Sèbe F, Bouix J, Boissy A. 2011 Assessment of sociability in farm animals: The use of arena test in lambs. *Applied Animal Behaviour Science* **135**, 57–62. (doi:10.1016/j.applanim.2011.09.004)

3. Thioulouse J, Dray S, Dufour A-B, Siberchicot A, Jombart T, Pavoine S. 2018 The dudi Class. In *Multivariate Analysis of Ecological Data with ade4* (eds J Thioulouse, S Dray, A-B Dufour, A Siberchicot, T Jombart, S Pavoine), pp. 29–51. New York, NY: Springer. (doi:10.1007/978-1-4939-8850-1_3)

4. Hazard D, Bouix J, Chassier M, Delval E, Foulquié D, Fassier T, Bourdillon Y, François D, Boissy A. 2016 Genotype by environment interactions for behavioral reactivity in sheep1. *Journal of Animal Science* **94**, 1459–1471. (doi:10.2527/jas.2015-0277)

5. Hazard D *et al.* 2022 118. Divergent genetic selections for social attractiveness or tolerance toward humans in sheep. In *Proceedings of 12th World Congress on Genetics Applied to Livestock Production (WCGALP)*, pp. 520–523. Rotterdam, the Netherlands: Wageningen Academic Publishers. (doi:10.3920/978-90-8686-940-4_118)
